# hnRNPUL1 ensures efficient Integrator-mediated cleavage of snRNAs and is mutated in amyotrophic lateral sclerosis

**DOI:** 10.1101/2023.07.22.550030

**Authors:** Ivaylo D. Yonchev, Carmen V. Apostol, Llywelyn Griffith, Ang Li, Larissa Butler, Helen Patrick, Elisa Aguilar-Martinez, Ashleigh G. R. Whelan, Caroline Evans, Mark J. Dickman, Johnathan Cooper-Knock, Pamela J. Shaw, Steven West, Ivana Barbaric, Ian M. Sudbery, Stuart A. Wilson

**Affiliations:** Sheffield Institute For Nucleic Acids (SInFoNiA) and School of Biosciences, The University of Sheffield, Sheffield, S10 2TN, UK; Department of Molecular and Cellular Biology, Baylor College of Medicine, Houston, TX 77030, USA; Department of Biochemistry and Molecular Pharmacology, Baylor College of Medicine, Houston, TX 77030, USA; UK Dementia Research Institute at King’s College London, London, WC1E 6BT, UK; Centre for Stem Cell Biology, School of Biosciences, The University of Sheffield, Sheffield, S10 2TN, UK; The University of Manchester, Faculty of Biology, Medicine and Health, Manchester, M13 9PT, UK; Department of Chemical and Biological Engineering, The University of Sheffield, Sheffield, S1 4LZ, UK; Sheffield Institute for Translational Neuroscience, The University of Sheffield, Sheffield, S10 2HQ, UK; Living Systems Institute, University of Exeter, Exeter, EX4 4QD, UK

## Abstract

Integrator cleaves nascent RNA, triggering RNA polymerase II transcription termination, but how cleavage is regulated is poorly understood. Here we show hnRNPUL1 ensures efficient Integrator-mediated cleavage of nascent RNA downstream of snRNA genes and, in the case of U2 snRNA, binds a terminal stem-loop involved in this process. In the nucleoplasm, hnRNPUL1 binds U4 snRNA and SART3 and enables efficient reformation of the U4:U6 di-snRNP for further rounds of pre-mRNA splicing. Sustained hnRNPUL1 loss leads to reduced levels of snRNAs, defects in histone mRNA 3′ end processing and loss of Cajal bodies. hnRNPUL1 binds RNA through multiple domains, including a globular central domain comprising tightly juxtaposed SPRY and dead polynucleotide kinase folds. This latter fold allows binding to 5′-monophosphorylated RNAs in a mutually exclusive manner with ATP binding and functions as an XRN2 antagonist when overexpressed. We identify a cohort of amyotrophic lateral sclerosis patients harbouring disruptive mutations in hnRNPUL1. SMN loss in spinal muscular atrophy and hnRNPUL1 loss both disrupt snRNP biogenesis, leading to motor neuron death, suggesting a common aetiology.

## Introduction

The heterogeneous nuclear ribonucleoproteins (hnRNPs) bind heterogeneous nuclear RNA, largely composed of pre-mRNA^1^. They play roles in transcription, RNA processing, stability, localisation and translation^2^. Mutations in hnRNP genes lead to multiple diseases, including frontotemporal dementia, amyotrophic lateral sclerosis and neurodevelopmental disorders^3–5^. hnRNPUL1 is one of the largest hnRNPs, sharing sequence homology with hnRNPU and hnRNPUL2. The well characterised hnRNPU plays roles in splicing through U2 snRNP control^6^ and chromatin retention of RNAs, including XIST^7^. It tethers RNA to chromatin via an N-terminal SAP domain, which is phosphorylated by Aurora B kinase during mitosis to drive RNA release^8^. All members of the U protein family have two central tightly juxtaposed domains: SPRY and a nucleotide triphosphate (NTP)-binding domain with Walker A and B motifs^9^, which in the case of hnRNPU allows ATP binding and regulates oligomerisation^10^. hnRNPU was previously assigned to a group of P-loop NTPases, including T4 polynucleotide kinase (PNK), by virtue of the 5-4-1-2-3 strand order in the central β-sheet of the NTP binding domain and presence of a helical “lid”^11^, which are conserved in hnRNPUL1. The SPRY/ATP-binding domain of hnRNPU also binds RNA^12^ and all U-family proteins have intrinsically disordered C-terminal domains (CTDs) with interspersed RGG motifs associated with RNA binding^13^.

In contrast to hnRNPU, the molecular characterisation of hnRNPUL1 has been more limited, although *in vivo* it has been implicated in several processes. hnRNPUL1 has been shown to play a role in DNA double-strand break (DSB) repair, where it is recruited to sites of damage via NBS1. As part of the MRE11-RAD50-NBS1 (MRN) complex, NBS1 promotes RNA polymerase II (PolII) transcription at DSBs, leading to production of dilncRNAs^14^. Further, hnRNPUL1 has been shown to work together with hnRNPUL2 to drive DNA end resection, ATR signalling and recruitment of BLM helicase, though its molecular role remains unclear^15^. hnRNPUL1 also plays a role in pre-mRNA splicing, specifically alternative splicing, though the molecular mechanisms are again not understood^16, 17^. It is also implicated in the repression of histone gene transcription in cell cycle-arrested cells through an interaction with the U7 snRNP^18^.

A large-scale analysis of chromatin binding identified hnRNPUL1 as the most highly enriched nuclear RNA-binding protein on snRNA genes, indicating a potential role in their biogenesis^19^. The 3′ processing of snRNAs involves the multisubunit Integrator complex, which is tightly associated with PolII^20^. The catalytic subunit, INTS11, drives 3′ cleavage, though how this process is regulated is unclear. Recent work revealed that Integrator plays a wider role in PolII transcription beyond snRNA genes, triggering early cleavage and termination on multiple PoII transcripts, including protein coding genes^21^.

Several hnRNPs are implicated in amyotrophic lateral sclerosis (ALS), including FUS, hnRNPA2B1 and hnRNPA1. A hallmark of these proteins is a prion-like domain that can self associate. When ranked on their likelihood of containing a prion-like domain, hnRNPUL1 is within the top ten human RNA-binding proteins^22, 23^. Moreover, hnRNPUL1 binds the ALS-causative FET family of proteins (FUS, TAF15, EWSR1)^24, 25^ and is sequestered by C9orf72 repeat expansion RNA, commonly associated with ALS^26^. Together, these data suggest hnRNPUL1 may play a role in ALS. In addition to a potential role in ALS, hnRNPUL1 has also been implicated in B-cell precursor acute lymphoblastic leukemia (ALL), where a chromosomal translocation fuses the hnRNPUL1 CTD to the DNA-binding domain of MEF2D to generate an aberrant transcription factor^27–29^.

Despite several studies investigating hnRNPUL1’s function, a molecular role remains elusive, partly because some studies reported inefficient knockdown of hnRNPUL1 by RNA interference^19^. To address its molecular role, we used auxin degron technology^30^, allowing efficient elimination of hnRNPUL1 from the cell. We reveal an essential role in snRNA biogenesis and recycling in the cell.

## Results

### hnRNPUL1 is a dead polynucleotide kinase

A distinguishing feature of the U protein family compared with other hnRNPs is the presence of a central NTP-binding domain. To understand the role of this domain in hnRNPUL1, we investigated its structure using an AlphaFold v2.0^31, 32^ model. This revealed a tight juxtaposition of the SPRY and NTP-binding domains, generating a single globular fold **(Figures 1A** and **1B)**. The confidence score reported for this fold was >90%, except for Thr^455^–Gln^486^ around a ligand binding pocket (discussed below), indicating a higher degree of local flexibility. The interface between the SPRY and NTP binding domains consisted of amino acid pairs with complementary electrostatic charge **(Figure S1A)**. The NTP-binding domain shares structural homology with the kinase domain of mammalian polynucleotide kinase phosphatase PNKP **(Figures 1C and S1C**, RMSD over 82 Cɑ pairs of 1.1 Å) and T4 PNK (RMSD across 59 Cɑ pairs of 0.9 Å). Based on experiments presented, we hereafter refer to the hnRNPUL1 NTP-binding domain as a dead polynucleotide kinase (dPNK).

**Figure 1.**
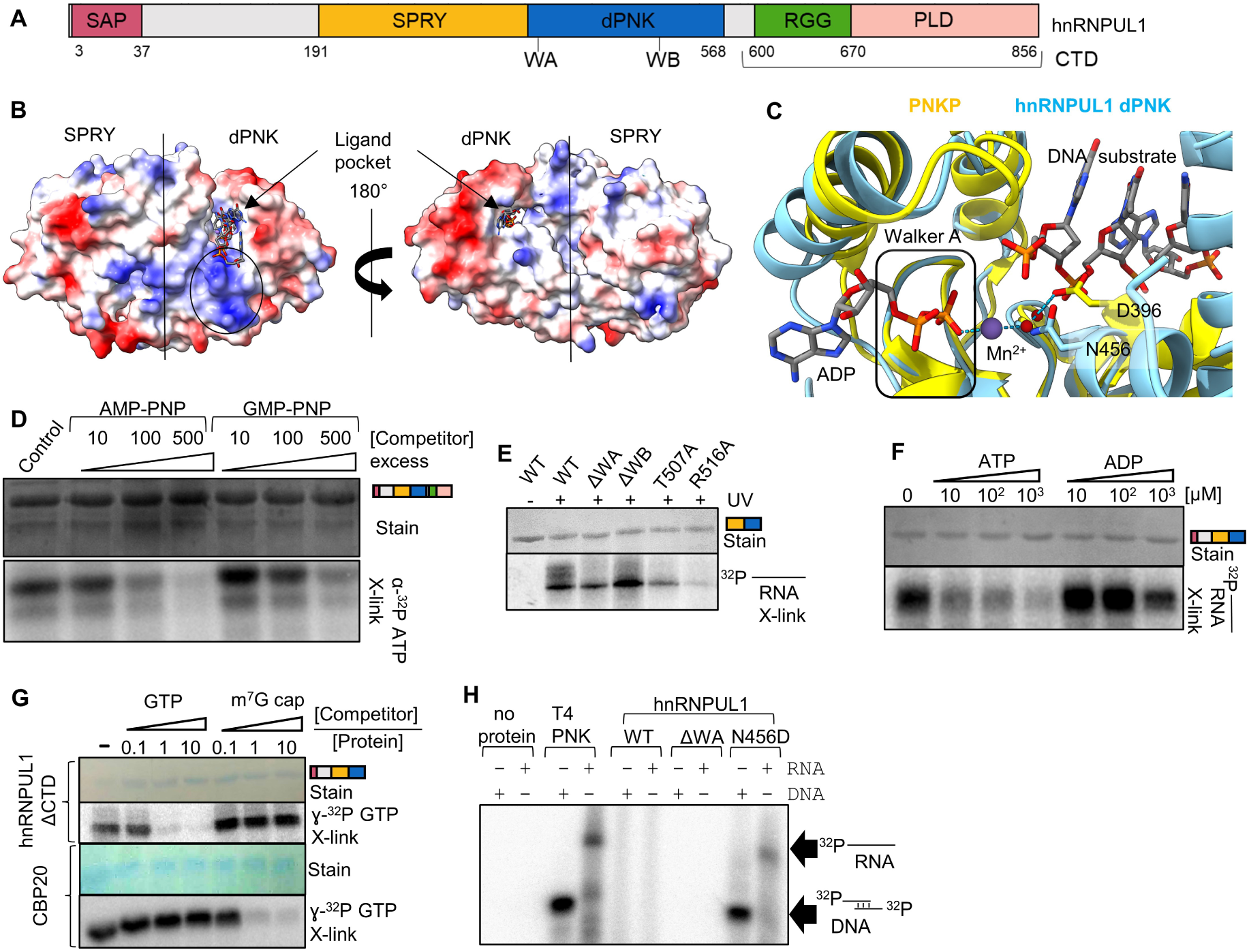
hnRNPUL1 dead polynucleotide kinase domain binds 5′ ends of non-capped RNA. **(A)** Schematics of hnRNPUL1 domain organisation. **(B)** AlphaFold v2.0 model of human hnRNPUL1 (Uniprot: Q9BUJ2) SPRY/dPNK “core” shown as surface charge. ATP and RNA ligands are docked into the predicted binding pocket based on homology with PNKP kinase domain. The basic patch leading into the binding pocket is encircled. **(C)** Structural comparison between PNKP liganded (yellow, PDB: 3ZVN) and hnRNPUL1 (blue) binding pockets. The sidechains of catalytic Asp396 (PNKP) and Asn456 (hnRNPUL1) are shown in stick form. Some regions are omitted for clarity. Dashed lines= hydrogen bonds, red spheres= oxygen atoms from water molecules. **(D)** Nucleotide competition of full-length FLAG-tagged hnRNPUL1 WT UV-crosslinked to ^32^P ɣ-ATP and varying amounts of non-hydrolysable ATP or GTP analogues. **(E)** UV-crosslinking of FLAG-tagged hnRNPUL1 SPRY/dPNK carrying different mutations to 5′ ^32^P-labelled RNA oligonucleotide. **(F)** UV-crosslinking of FLAG-tagged hnRNPUL1 ΔCTD truncation to 5′ ^32^P-labelled RNA oligonucleotide and varying amounts of ATP or ADP. **(G)** Competition assay of FLAG-tagged hnRNPUL1 ΔCTD (or 6xHis-tagged CBP20 as a positive control) UV-crosslinked to ^32^P ɣ-GTP and varying amounts of unlabelled GTP or m^7^G cap analogue. **(H)** Kinase assay using full-length FLAG-tagged hnRNPUL1 variants, ^32^P ɣ-ATP and RNA or dsDNA oligonucleotide substrates with free 5′ ends and 5′ overhangs. T4 PNK was used as a positive control.

Using UV cross-linking, we found that hnRNPUL1 bound ATP via its dPNK domain and not through non-specific electrostatic interactions with the RGG box **(Figure S1B).** A mutation in the Walker A motif (ΔWA=^428^GLPAA<u>AAA</u>^435^) reduced ATP binding, whereas a Walker B mutation (ΔWB= ^502^YIL<u>A</u>^505^) enhanced it **(Figure S1B)**. The intrinsic tryptophan fluorescence of the hnRNPUL1 core altered upon ATP binding, indicative of a conformational change **(Figure S1C)**, a feature common to PNKP upon binding its ligands^33^. Binding constants derived for WT, ΔWA and ΔWB proteins support the UV-crosslinking results: Kd_WT_= 164 ± 22 nM, Kd_WA_= 430 ± 247 nM and Kd_ΔWB_= 19 ± 8.5 nM. Trp^477^ and Trp^437^ are located close to the binding site and likely contribute to the fluorescent signal quenching. Moreover, Trp^477^ is found in a region with a predicted high degree of flexibility according to the AlphaFold model (<70% confidence), further implicating this region in the protein’s ligand-induced conformational change. In a competition assay, non-hydrolysable analogs of both ATP and GTP displaced ^32^P-labelled ATP from hnRNPUL1, which is consistent with the ability of T4 PNK to use a range of NTPs as phosphate donors **(Figure 1D).**

Since polynucleotide kinases bind nucleic acids, we next tested whether the isolated SPRY/dPNK domain of hnRNPUL1 could bind RNA. The wild-type (WT) SPRY/dPNK domain bound a 5′-monophosphorylated (monoPi) RNA oligonucleotide, whilst ΔWA displayed marginally reduced binding and ΔWB a slightly enhanced binding **(Figure 1E)**. The ability of ΔWB to bind ATP **(Figure S1B)** and RNA better may be explained by the creation of a less acidic phosphate-binding pocket **(Figure S1D)**. We noted a basic track leading into the ligand-binding pocket of hnRNPUL1 **(Figure 1B)** equivalent to a DNA-binding site in PNKP **(Figure S1D)**. Mutation of two residues within this track (T507 and R516) disrupted RNA binding **(Figure 1E)**, indicating conservation of nucleic acid-binding sites between hnRNPUL1 and PNKP (T423 and R432). Further supporting the SPRY/dPNK cores of hnRNPUL1 and hnRNPU as RNA-binding domains, peptides from these regions were recently identified in global mass spectrometry studies of RNA-binding proteins^12, 34–36^ **(Figure S1E)**. Additional RNA-binding peptides were mapped in U-family proteins to the SAP and the well characterised RGG RNA-binding domain^13, 37^, indicating they make contact with RNA through multiple domains *in vivo*.

To explore the interplay between ATP and RNA binding to the hnRNPUL1 dPNK domain, we assessed binding to a 5′-monoPi RNA in the presence of ADP and ATP **(Figure 1F)**. ATP effectively outcompeted a 5′-monoPi RNA, whereas ADP did not, indicating the pocket can accommodate up to 3 phosphates and that nucleotide triphosphates may play a role in RNA ligand turnover. The arrangement of ADP and a 5′-monoPi RNA accessing the binding pocket from opposite sides was reminiscent of the inverted 5′ m^7^G cap structure of mRNA. Therefore, we investigated whether a triphosphate-linked cap analogue could access the substrate pocket. Unlabelled GTP was an effective competitor for radioactive GTP binding, whereas an m^7^G cap analog was not **(Figure 1G)**. For comparison, m^7^G displaced GTP from the cap-binding protein CBP20, indicating that hnRNPUL1 is unlikely to accommodate the 5′ cap structure of an mRNA in its dPNK pocket.

We noted a key difference in the binding pockets of hnRNPUL1 and PNKP: the catalytic D396 of PNKP is an Asn at the equivalent position in hnRNPUL1. A D396N mutation in PNKP abolishes its kinase activity^38^. Therefore, we speculated that reversing N456 to Asp may restore a polynucleotide kinase activity to hnRNPUL1. Remarkably, in an ATPase assay, the N456D mutant, but not WT, generated ∼50% more ADP over background, while addition of a 5′-unphosphorylated RNA oligonucleotide stimulated ATP hydrolysis by >100% **(Figure S1F)**. Strikingly, a kinase assay revealed that hnRNPUL1 N456D and T4 PNK were able to phosphorylate both RNA and dsDNA substrates with free 5′ ends (and overhangs), whereas hnRNPUL1 WT and ΔWA were not **(Figure 1H)**. hnRNPU also contains an Asn at the equivalent position (N512) in its dPNK domain, whereas hnRNPUL2 possesses an acidic Glu residue (E487). However, a kinase assay with hnRNPUL1 N456D, hnRNPU N512D and hnRNPUL2 WT revealed that the reactivation of the nucleic acid kinase function was specific to the hnRNPUL1 mutant and not a common feature of the U-family achievable with a point mutation **(Figure S1G)**, despite their extensive predicted structural homology. Together, these findings demonstrate that, *in vitro,* hnRNPUL1 is a dead polynucleotide kinase which has retained the ability to bind nucleotide triphosphates and RNA, validating the structural prediction.

### hnRNPUL1 is required for U4-U6 di-snRNP formation

Having established the dPNK domain of hnRNPUL1 could bind ATP and ligand binding induced a conformational change, we investigated the role of this domain initially using mass spectrometry to identify binding partners. We generated HEK293 T-Rex FLP-In cell lines for FLAG-tagged WT, ΔWA and a mutant lacking the CTD and determined conditions for their expression at levels close to endogenous hnRNPUL1 **(Figure S2A)**. hnRNPUL1 binding partners were identified for both WT and mutant proteins in the presence and absence of RNAse A **(Table S1)**. hnRNPUL1 interactors included RNA-binding proteins such as ribosomal proteins, core snRNP components, and other proteins linked to various stages of RNA metabolism, including RNA turnover **(Figure S2B)**. Several additional spliceosomal and snRNP biogenesis components were identified as RNA-dependent hnRNPUL1 interactors **(Table S1)**. Interestingly, we observed an altered protein interactome in the case of ΔWA, with enhanced RNA-independent interactions with several binding partners, including the FET protein family. For this reason, we investigated the cellular distribution of hnRNPUL1 WT and ΔWA in these cell lines and observed a contrasting localisation, with equal levels of WT protein in the nucleoplasm and chromatin compartments and >80% ΔWA mutant on chromatin **(Figure 2A)**. The entrapment of the ΔWA mutant in the chromatin environment may result from reduced solubility or failure to induce a conformational change upon nucleotide triphosphate binding and release of an RNA substrate. Nevertheless, the altered distribution reveals hnRNPUL1 associates with distinct groups of proteins in each environment.

**Figure 2:**
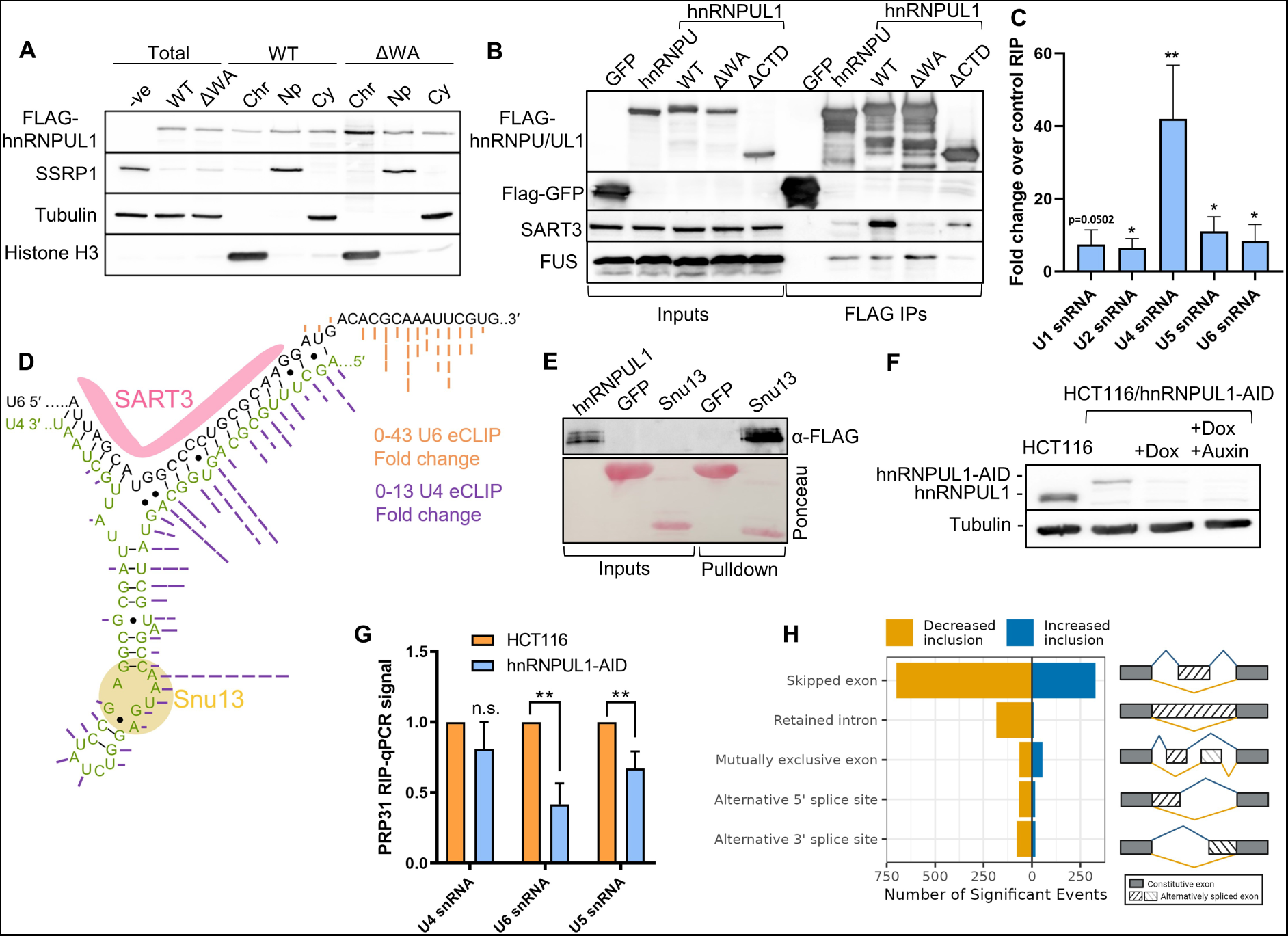
hnRNPUL1 is required for efficient U4/U6 di-snRNP formation and normal splicing. **(A)** Fractionation of hnRNPUL1 WT and ΔWA using stable FlpIn cell lines. **(B)** Co-IP of FLAG-tagged hnRNPU and hnRNPUL1 variants showing interaction with SART3 identified by mass spectrometry. ΔWA and ΔCTD mutations abolish or reduce interactions, respectively. **(C)** RIP-qPCR showing hnRNPUL1 enrichment over snRNAs (n = 3). Immunoprecipitated RNAs were measured as a percentage of input, then expressed as fold change over a mock antibody RNA-IP. **(D)** Schematic of U4/U6 snRNA base pairing highlighting hnRNPUL1 eCLiP signal (dashed lines) and positions of U4 snRNP factor Snu13 and U6 snRNP factor SART3. Black lines= canonical pairs, black dots= non-canonical pairs. **(E)** Pulldowns of FLAG-tagged hnRNPUL1 (prey) using 6xHis-tagged Snu13 (bait) and 6xHis-tagged GFP (negative control). **(F)** hnRNPUL1 levels before and after treatment of hnRNPUL1-AID cells. **(G)** RIP-qPCR of U4 snRNP factor PRP31 in HCT116 and hnRNPUL1-AID cells (n = 3). Immunoprecipitated RNAs were measured as a percentage of input, then expressed as fold change over the HCT116 RNA-IP. **(H)** Alternative splicing events in hnRNPUL1-AID cells. Bar charts show relative mean values from at least three independent experiments with error bars showing s.d. All statistical analyses were performed using two-tailed t-tests. n.s. = not significant, *p<0.05, **p<0.01.

Comparison of the WT and ΔWA interactomes revealed a striking loss of the U6 snRNP-specific factor SART3 **(Figure S2C)**. This protein shuttles between the nucleoplasm and Cajal bodies, promoting formation of the U4-U6 di-snRNP, which subsequently associates with U5 snRNP before entering the splicing reaction. U4 and U6 snRNA annealing is disrupted during splicing, and each component is released. SART3 facilitates recycling of these snRNPs for further splicing^39^. Co-IP confirmed a robust interaction between hnRNPUL1 and SART3 **(Figure 2B)**. Notably, ΔWA displayed a greatly reduced association, consistent with the MS data **(Figure S2C)**. This is in direct contrast with the enhanced binding observed between ΔWA and chromatin-associated interactors, such as FUS **(Figure 2B, Table S1),** suggesting the interaction between hnRNPUL1 and SART3 occurs in the nucleoplasm. Cellular fractionation confirmed SART3 is primarily nucleoplasmic and cytoplasmic but absent from chromatin **(Figure S2D)**. Thus, the chromatin-biassed distribution of ΔWA most likely prevents its association with SART3.

Given the prominent interaction between hnRNPUL1 and SART3, we performed RNA-Immunoprecipitation (RIP) qRT-PCR using an antibody to endogenous hnRNPUL1 to investigate hnRNPUL1:snRNA interactions. Interestingly, U4 snRNA, followed by U5 and U6 snRNAs, displayed the strongest interaction with hnRNPUL1 **(Figure 2C)**, though this interaction is unlikely to involve direct docking within the P-loop region of the dPNK domain, since this site is unable to accommodate the 5′ cap found on mature U4 snRNA **(Figure 1G)**. We analysed ENCODE enhanced crosslinking with immunoprecipitation (eCLIP) hnRNPUL1 data for signal over the U6 and U4 snRNAs^40^ **(Figure 2D)**. hnRNPUL1 bound U6 snRNA downstream of the SART3 binding site and the 5′ end and 5′ stem loop of U4 snRNA, around the binding site for the U4-specific factor Snu13 (also called NHP2L1 or 15.5K). This protein is a recruitment platform for additional U4 snRNP factors, such as Prp31 and CypH/Prp3/Prp4^41^. We detected a direct and specific interaction between Snu13 and hnRNPUL1 by pulldown **(Figure 2E)**. Next, we tested a series of hnRNPUL1 mutants for association with Snu13 and found that deletion of the SPRY domain was sufficient to disrupt Snu13 binding, highlighting an important role for this domain in Snu13 interactions **(Figure S2E)**.

To address the role of hnRNPUL1 function in U4-U6 di-snRNP formation and subsequent U4-U6/U5 tri-snRNP assembly, we tagged the gene with an auxin-inducible degron in HCT116 cells harbouring a doxycycline inducible TIR1 gene, enabling depletion of hnRNPUL1 with doxycycline and auxin (Dox/Aux) treatment **(Figure 2F)**. Despite some basal degradation of hnRNPUL1 in the absence of treatment, cell survival was not affected, yet elimination of hnRNPUL1 with Dox/Aux inhibited cell proliferation **(Figure S2F)**. We then assessed U4-U6/U5 tri-snRNP assembly using RIPs with the U4 snRNP-specific factor Prp31 and found that depletion of hnRNPUL1 disrupted di- and tri-snRNP formation **(Figure 2G)**. Furthermore, analysis of chromatin-associated RNA-seq data from our hnRNPUL1-AID cells revealed over a thousand significantly altered splicing events triggered by loss of hnRNPUL1 **(Figure 2H)**. Reanalysis of ENCODE RNAseq data following knockdown of hnRNPUL1 revealed similar effects^40^, indicating that hnRNPUL1 loss disrupts splicing fidelity **(Figure S2G)**. Together these data reveal that in the nucleoplasm, hnRNPUL1 functions together with SART3 to ensure efficient assembly and/or recycling of the U4-U6/U5 tri-snRNP.

### hnRNPUL1 is required for Cajal body formation

The recycling of the U4-U6 di-snRNP occurs in Cajal bodies (CB), which are nuclear membraneless organelles that direct multiple steps in snRNP biogenesis and recycling^42^. CBs are enriched for SART3 and the archetypal CB marker protein coilin^42, 43^, as well as multiple other RNAs including snoRNAs and scaRNAs. To determine whether hnRNPUL1 bound other CB-associated RNAs, we analysed eCLIP datasets^40^ and categorised RNA species based on biotype. hnRNPUL1 showed enrichment over scaRNA, snoRNA and snRNA categories, in contrast to other hnRNPs **(Figures 3A and S3A)**. There was an exceptional enrichment of hnRNPUL1 on U7 snRNA, consistent with a previous report of its role in regulating histone transcription via U7 snRNA^18^. The eCLIP enrichment pattern of hnRNPUL1 was reminiscent of coilin’s RNA interactome^42^. Furthermore, hnRNPUL1 and coilin ChIP-seq datasets revealed coincident hnRNPUL1 binding at coilin peaks **(Figures S3B and S3C)**. We therefore examined the role of hnRNPUL1 in CB formation. Despite reduced levels of hnRNPUL1 due to basal degradation in hnRNPUL1-AID cells, there was no reduction in the levels of CBs. However, following Dox/Aux addition and complete hnRNPUL1 knockdown, we saw a dramatic reduction in CB numbers **(Figures 3B and 3C)**. The depletion of hnRNPUL1 also resulted in the loss of SMN-containing nuclear Gems, which frequently co-localise with CBs^44^ **(Figure 3B)**, and disruption of the coilin interactome including SMN, SART3 and U1 snRNP components **(Figure 3D)**.

**Figure 3:**
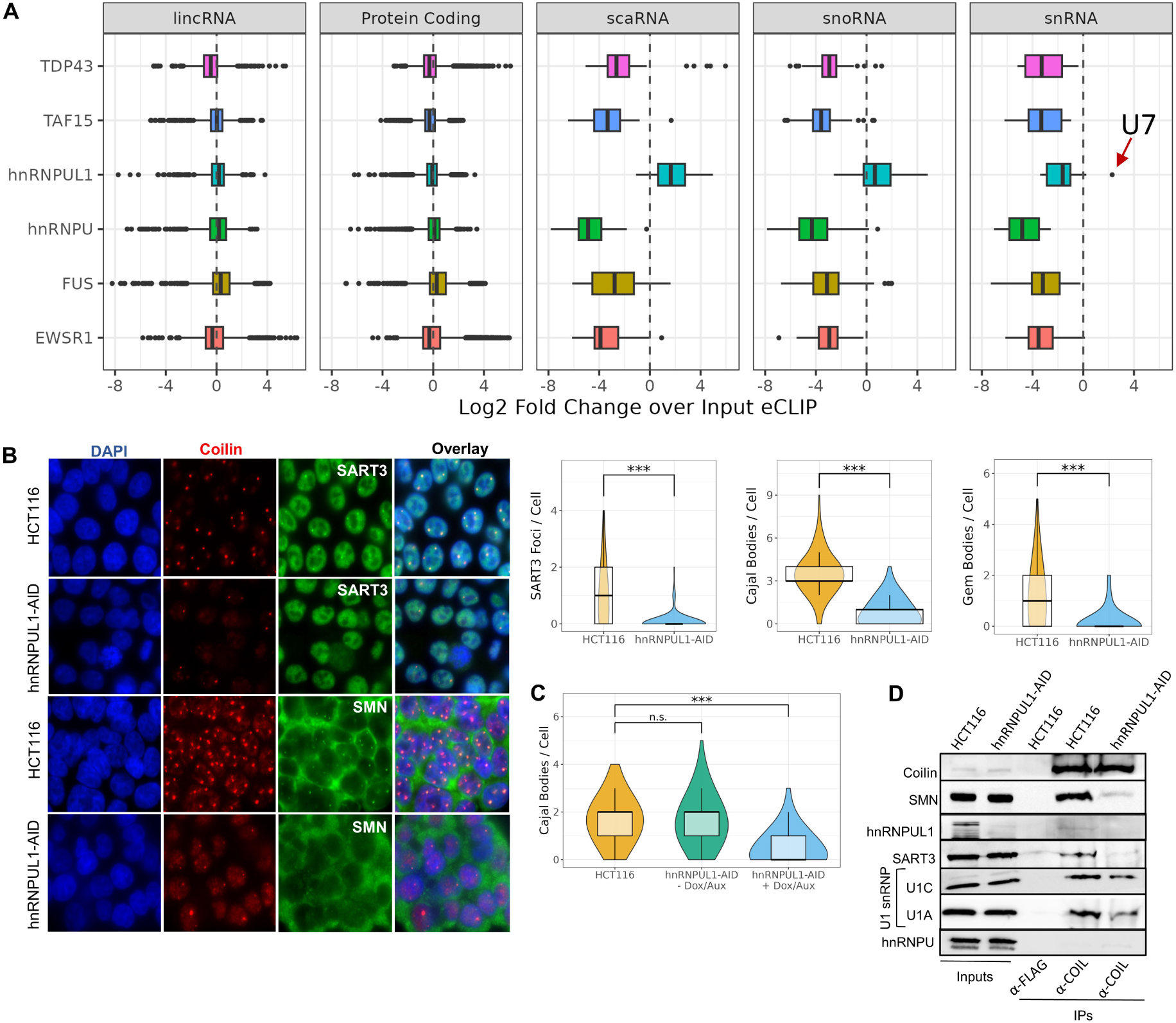
hnRNPUL1 associates with the coilin interactome and is required for Cajal body formation. **(A)** Log2 Fold Change DESeq2 normalized eCLIP signal over input for select RNA biotypes. **(B)** Immunofluorescence images and violin and box plot quantification of coilin (Cajal Body), SART3 and SMN (Gem Body) foci per cell. **(C)** Violin and box plot quantification of Cajal Bodies showing unaffected numbers in hnRNPUL1-AID cells before drug treatment. Violin and box plots show the median and 25-75^th^ percentile values from two independent experiments for public eCLIP data and three independent experiments for nuclear body quantification, containing 60-80 cells counted for each replicate. All statistical analyses were performed using Welch’s t-test. n.s. = not significant, ***p<0.001. **(D)** Co-IP of coilin interactome in hnRNPUL1-AID cells.

### hnRNPUL1 CTD binds PolII CTD

Having identified a nucleoplasmic role for hnRNPUL1, we turned our attention to its role in the chromatin environment. We were interested in establishing the localisation of ΔWB, therefore we expressed this mutant, along with WT and ΔWA, in the hnRNPUL1-AID cell line and depleted the degron-tagged endogenous protein. This experiment established that ΔWB was also trapped in the chromatin fraction **(Figure S4A)** and the localisation patterns of WT and ΔWA were similar at both high and low expression levels, indicating disrupted localisation is not simply a consequence of overexpression **(Figure S4B)**. Therefore, mutations which either disrupt NTP binding (ΔWA) or enhance it together with RNA binding (ΔWB) alter the distribution of hnRNPUL1. Next, we tested the mode of chromatin association of hnRNPUL1 WT, ΔWA and ΔWB by treating chromatin fractions with RNase A and subsequently DNase and monitoring protein release **(Figure 4A)**. Most of the WT protein was released by RNase, indicating it is largely engaged with chromatin via RNA. However, ΔWA and ΔWB were biassed towards DNase release, indicating a fundamentally different association with chromatin. Together, these data imply that either protein-protein or protein-DNA interactions might also be perturbed by these mutations in the nucleotide triphosphate-binding site, consistent with our MS results. Strikingly, crosslinking immunoprecipitation (CLIP) analysis **(Figure 4B)** revealed that both ΔWA and ΔWB crosslinked with more RNA than WT protein *in vivo*, as well as *in vitro* **(Figure S4C)**. Therefore, the failure to be released by RNase was not due to a lack of RNA binding. The dramatic increase in RNA binding *in vivo* observed with ΔWA, which is unable to bind ATP **(Figure S1B)**, is consistent with the *in vitro* observation that ATP competes for RNA binding **(Figure 1F)**. The increased RNA binding observed with ΔWB is again consistent with the *in vitro* observation that this mutation creates a more favourable RNA binding pocket **(Figure 1E)**. However, we cannot exclude the possibility that the trapping of these mutant proteins in the RNA-rich chromatin environment also contributes to the enhanced RNA crosslinking *in vivo*.

**Figure 4:**
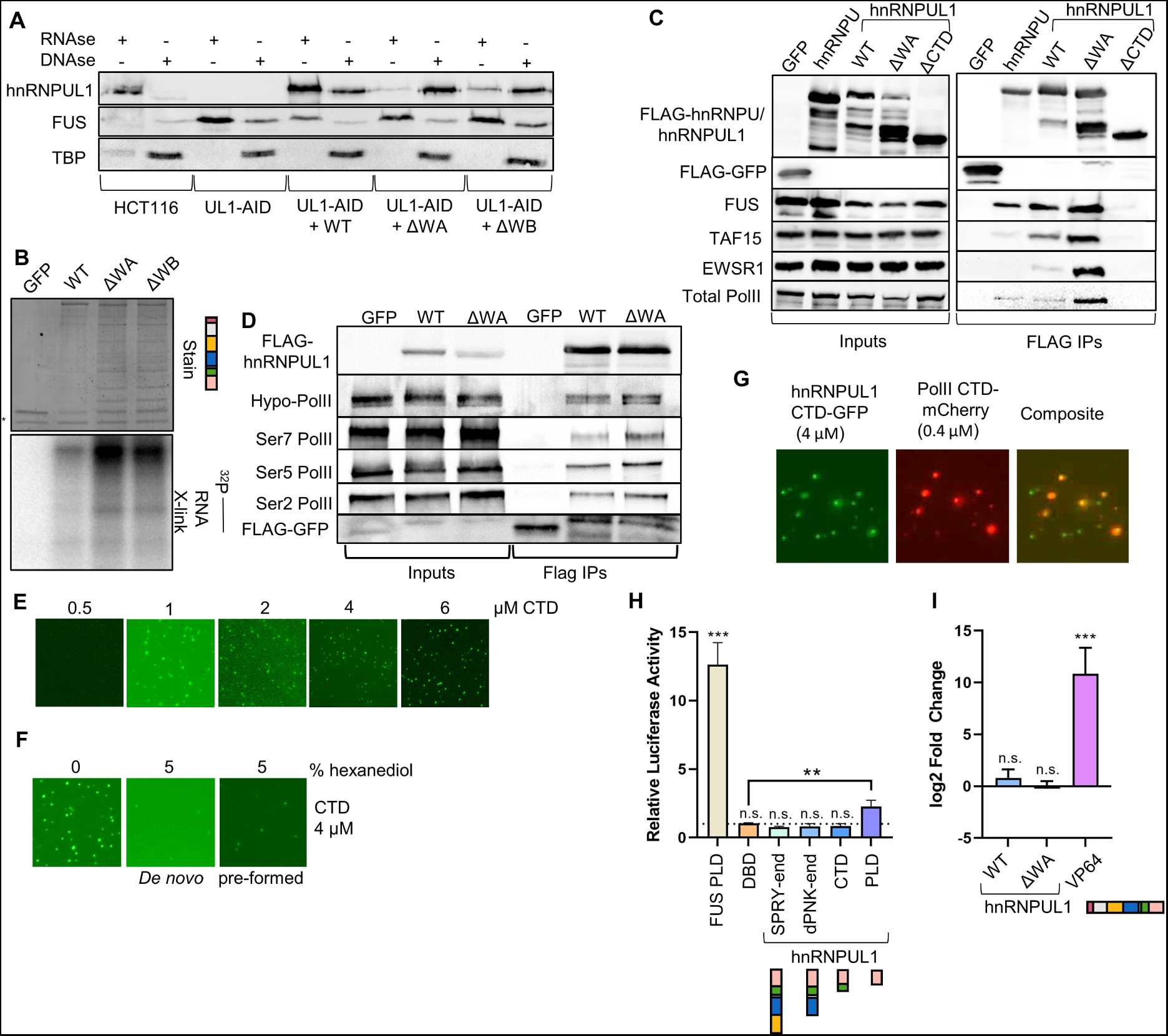
hnRNPUL1 CTD associates with RNA polymerase II CTD. **(A)** Chromatin release of WT hnRNPUL1 in HCT116 cells and FLAG-hnRNPUL1 WT/ΔWA/ΔWB expressed in hnRNPUL1-AID cells. Protein was first released using RNAse treatment and this was followed by DNAse mediated release. **(B)** *In vivo* crosslink-IP (CLIP) of FLAG-tagged hnRNPUL1 WT/ΔWA/ΔWB expressed in HEK293T, followed by FLAG-IP, partial RNase A digestion and T4 PNK end-labelling of the RNA with ^32^P ɣ-ATP. *= T4 PNK. **(C)** Co-IP of hnRNPU and hnRNPUL1 variants identifying hnRNPUL1 CTD as a crucial mediator of interactions with FET proteins and PolII. **(D)** hnRNPUL1 WT/ΔWA co-IP assessing interaction with PolII and its CTD phospho-isoforms. **(E)** LLPS assays of hnRNPUL1 CTD expressed with N-terminal GFP and C-terminal 6xHis tags at the specified concentrations. **(F)** LLPS assays of hnRNPUL1 CTD (4 µM) with 5% 1,6-hexanediol added at the beginning of the incubation (de novo) or after 1 hr LLPS incubation (pre-formed). **(G)** Co-localisation of GFP-tagged hnRNPUL1 CTD and mCherry-tagged PolII CTD within phase separated droplets. **(H)** Dual luciferase reporter assay measuring transcription from a GAL4-driven promoter after transfection of GAL4-fused FUS and hnRNPUL1 constructs (n = 4). DBD= DNA-binding domain. **(I)** qRT-PCR measurement of the change in RNA production, relative to 18S rRNA, in HEK293T cells, after hnRNPUL1 and VP64 tethering to ASCL1 gene over dCas9 + gRNA background control (n = 3). The bar charts show relative mean values from at least three independent experiments, with the error bars showing s.d. All statistical analyses were performed using two-tailed t-tests. n.s. = not significant, **p<0.01, ***p<0.001.

An earlier MS study identified hnRNPUL1 as a prominent PolII interactor^45^. Given its ability to bind chromatin, we considered that the ΔWA mutation might enhance its interaction with PolII, as it does with FUS **(Figure 2B)**, and this may be partly responsible for the ability of DNAse, but not RNAse, to release ΔWA from chromatin. Co-IP analysis revealed that ΔWA bound not only PolII, but also three FET proteins, better than the wild-type protein and revealed an essential role for hnRNPUL1 CTD for these interactions **(Figure 4C)**. hnRNPUL1 interacted with Ser2,5 and 7-phosphorylated PolII, with ΔWA displaying enhanced binding to all phosphoisoforms, indicating it associates with actively-transcribing PolII **(Figure 4D)**. We identified minimal regions for interaction between hnRNPUL1 and PolII using the PolII CTD as bait. Only full-length hnRNPUL1 and the isolated CTD came down with PolII CTD in an RNA-independent manner, suggesting that the interaction requires their CTDs **(Figure S4D)**.

To further investigate the nature of the CTD of hnRNPUL1, we analysed its sequence using the PLAAC platform^46, 47^ (Prion-Like Amino Acid Composition), which identified a prion-like domain (PLD, **Figure S4E)**. Moreover, the CTD exhibits features characteristic of phase separating proteins, such as an unstructured nature, low amino acid diversity, Arg-Tyr bias in the RGG box and PLD and (Ser/Gly)Tyr(Ser/Gly) prion-like repeats^48, 49^. Therefore, we investigated its phase behaviour. A GFP-CTD fusion formed protein droplets in physiological sodium conditions (100-150 mM) and in a concentration-dependent manner starting around 1 µM protein **(Figure 4E)**. Pre-formed droplets were dissolved by 1,6-hexanediol, which also prevented new droplet formation, indicating the droplets likely resulted from liquid-liquid phase separation (LLPS), not protein aggregation **(Figure 4F)**. The hnRNPUL1 CTD droplets concentrated PolII CTD from the surrounding solution, underscoring the importance of the hnRNPUL1 and PolII CTDs for their interaction **(Figure 4G)**. Notably, a similar PolII CTD construct was reported to fail to phase separate on its own^50^, while in our hands, it formed protein aggregates of irregular shapes and sizes **(Figure S4G)**, further suggesting that the hnRNPUL1 CTD promotes concentration of PolII CTD into round droplets.

Given the hnRNPUL1 CTD-PolII CTD interaction and the previously reported transcriptional activity of MEF2D-hnRNPUL1 CTD fusions^27–29^ **(Figure S4F)**, we tested whether hnRNPUL1 might function as a transcription factor. We fused hnRNPUL1 truncations to the GAL4 DNA-binding domain and monitored gene expression with a dual luciferase reporter **(Figure 4H)**. Only the hnRNPUL1 PLD increased transcription of the reporter, whilst less extensive hnRNPUL1 truncations were inactive. Concomitantly, tethering 24 copies of full-length WT or ΔWA to a dead Cas9 at the ASCL1 gene promoter failed to activate transcription above background, unlike the VP64 positive control **(Figure 4I)**. Thus, intact hnRNPUL1 is unlikely to drive PolII transcription from protein-coding gene promoters, but its CTD displays a gain-of-function transcription factor activity when expressed as a fusion with a heterologous DNA binding domain, such as the MEF2D DNA-binding domain in ALL. Taken together, our results demonstrate that the hnRNPUL1 CTD is required for interaction with multiple proteins, including PolII, and the dPNK domain enables its dissociation from chromatin-associated RNA and certain interacting proteins.

### hnRNPUL1 is necessary for snRNA 3′ processing and transcription termination

The ability of hnRNPUL1 to load onto actively transcribing PolII CTD via its own CTD suggested it may play a role in co-transcriptional processing of target RNAs. snRNAs were a strong candidate for such a function, given the strong enrichment of hnRNPUL1 ChIP signal at snRNA genes^19^. Using the hnRNPUL1-AID degron line, we performed small non-coding RNA sequencing (sRNA-seq; size range 50-500 bases), which revealed downregulation of snRNAs upon hnRNPUL1 depletion **(Figure 5A)**. PolII ChIP showed decreased PolII occupancy at snRNA gene bodies following hnRNPUL1 loss **(Figure 5B)**, indicating reduced transcription. This decrease was commensurate with the decrease in snRNA levels observed in our sRNA-seq and through qRT-PCR **(Figure S5A)**. The hnRNPUL1 ChIP-seq binding profile over snRNA genes showed greatest enrichment downstream of the 3′ RNA cleavage site **(Figures 5C and S5B)**. Thus, we investigated whether snRNAs underwent hnRNPUL1-dependent cleavage and termination. Metagene analysis of chromatin RNA-seq data over primary PolII-transcribed snRNA loci revealed decreased signal over the gene body, followed by increased density downstream of the cleavage site following hnRNPUL1 loss **(Figure 5D, S5C)**. We also used qRT-PCR with primers spanning the 3′ cleavage site to confirm there was a 3′ cleavage defect for snRNAs following hnRNPUL1 depletion **(Figure S5D)**. Similar snRNA metagene profiles have been reported following depletion of the catalytic subunit of the Integrator complex INTS11^51^. To further enrich the nascently transcribed subpopulation of snRNAs, we performed transient transcriptome sequencing with chemical RNA fragmentation (TTchem-seq), which showed increased signal beyond the 3′ end of snRNA genes within a 5 minute 4SU pulse-chase **(Figures 5E, S5C and S5E)**. Interestingly, we also saw increased nascent RNA signal across snRNA gene bodies. Both of these effects match those observed using TT-seq following acute depletion of Ints11^21^. Failure to cleave snRNAs at the 3′ end is expected to lead to a PolII termination defect and PolII ChIP across the U1 snRNA locus confirmed increased occupancy 550 bp downstream of the snRNA gene body relative to the proximal sequence element following hnRNPUL1 loss, consistent with a termination defect **(Figure 5F)**.

**Figure 5.**
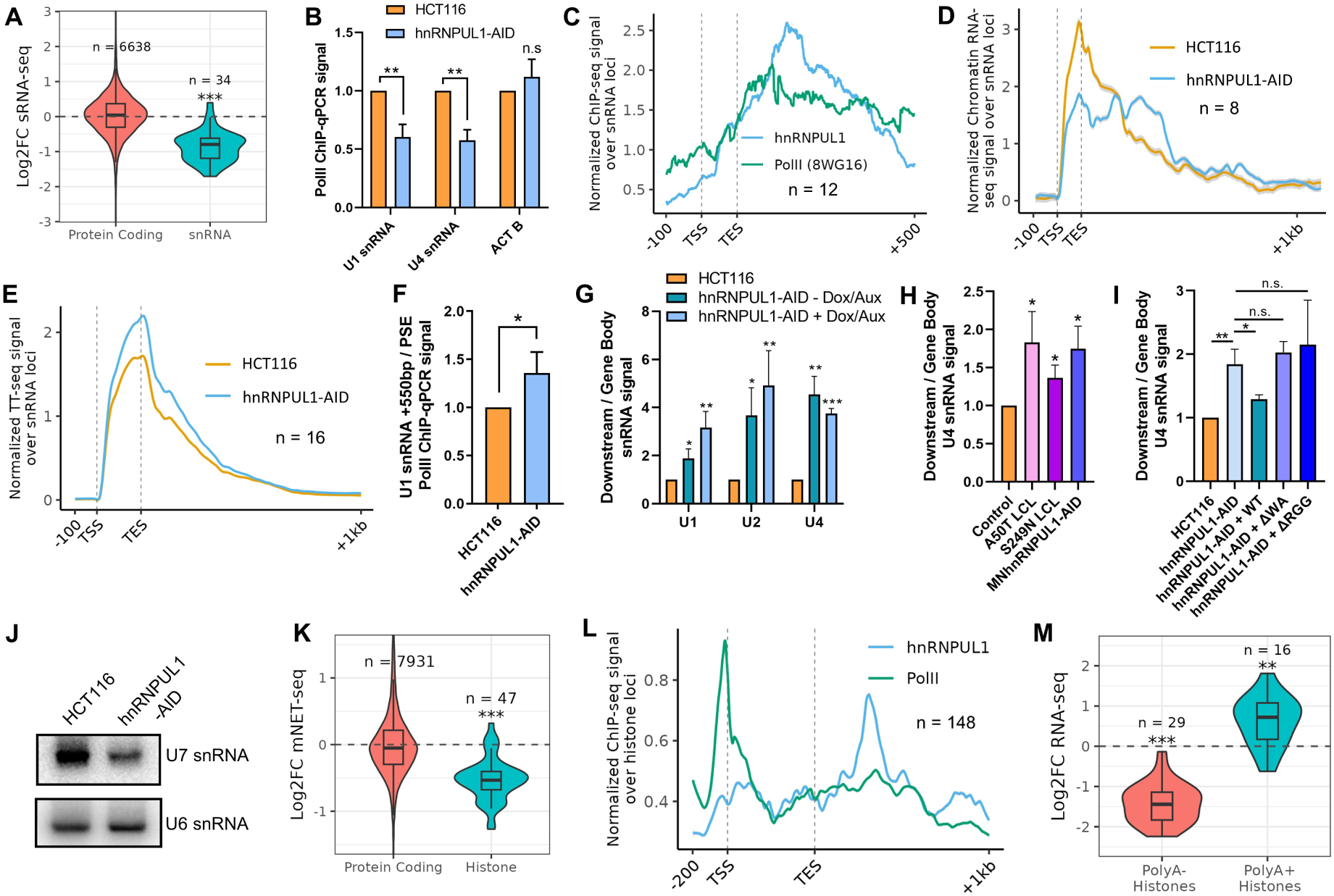
Loss of hnRNPUL1 leads to cleavage and transcription termination defects on snRNAs and histone mRNAs. **(A)** snRNA levels in hnRNPUL1 cells measured through sRNA-seq. Although sRNAs were enriched in these datasets, counts from other biotypes, such as protein coding RNAs, were present and used to aid sample normalisation. All snRNA annotations containing mapped reads, including variants, were included in the analysis. **(B)** PolII ChIP-qPCR over snRNA gene loci (n = 3). Immunoprecipitated chromatin was measured as a percentage of input, then expressed as fold change over the HCT116 ChIP. **(C)** Metagene of hnRNPUL1 and PolII ChIP-seq fold change over input signal distribution over non-variant PolII-transcribed snRNA gene annotations, containing mapped reads, and their downstream region. TSS = Transcription Start Site, TES = Transcript End Site. **(D)** Metagene of HCT116 and hnRNPUL1-AID chromatin RNA-seq signal distribution over PolII-transcribed snRNA loci. Loci containing >50,000 counts across each 1.2kb region were considered mature snRNAs and were removed from the final plot. **(E)** Metagene of DESeq2-normalized TT-seq signal, summed across all non-variant PolII-transcribed snRNA gene annotations and their downstream region. **(F)** PolII ChIP-qPCR showing the ratio between the region 550 bp downstream of U1 and its proximal sequence element (PSE) in hnRNPUL1-AID cells relative to HCT116 (n = 3). **(G)** Levels of 3′-extended snRNAs measured through the ratio between downstream and gene body-amplified snRNA levels in untreated and Dox/Aux-treated hnRNPUL1-AID cells (n = 3). **(H)** 3′-extended U4 snRNA levels in A50T and S249N Lymphoblastoid and untreated motor neuron-differentiated hnRNPUL1-AID stem cells, measured as in (G) (n = 3). A LCL generated from a healthy individual of similar age was used as the control for A50T and S249N LCL conditions, while motor neuron-differentiated stem cells not containing the hnRNPUL1-AID were used as the control for the MNhnRNPUL1-AID condition. **(I)** 3′-extended U4 snRNA levels in untreated hnRNPUL1-AID cells and Sleeping Beauty transposase-complemented hnRNPUL1-AID cells, measured as in (G) (n = 3). **(J)** Northern Blot of U7 snRNA levels in HCT116 and hnRNPUL1-AID cells. U6 snRNA is shown as a loading control. **(K)** PolII density over Protein Coding and replication-dependent Histone genes measured through mNET-seq. **(L)** Metagene of hnRNPUL1 and PolII ChIP-seq fold change over input signal, summed across all Histone genes and their downstream region. **(M)** Levels of replication-dependent histones in nuclear (polyA-) and (polyA+) RNA-seq data. Bar charts show relative mean values from at least three independent experiments with error bars showing s.d. Violin and box plots show median and 25-75^th^ percentile values from three independent experiments. Bar plot statistical analyses were performed using two-tailed t-tests, while violin and box plot statistical analyses were performed using Welch’s t-test. n.s. = not significant, *p<0.05, **p<0.01, ***p<0.001.

### hnRNPUL1 is mutated in ALS patients

Pre-mRNA splicing, CBs and SMN-containing Gem structures are all commonly disrupted in ALS and spinal muscular atrophy (SMA)^44, 52–54^. In light of this, and the association of hnRNPUL1 with other ALS-causing proteins, C9orf72 repeats and its prion-like characteristics, we screened two cohorts of familial (n=1,022, ALS variant server) and sporadic (n=4,366)^55^ ALS patients for pathogenic coding mutations within hnRNPUL1. We defined “rare” based on Minor Allele Frequency < 1% in population databases^56^ and “pathogenic” based on Combined Annotation Dependent Depletion score >10^57^. Multiple heterozygous variants identified using these criteria are shown in **Table S2**. Notably, mutation R541X involved a severe truncation, suggesting loss of hnRNPUL1 function may be associated with ALS.

To examine the effects of ALS mutations in hnRNPUL1, we obtained patient-derived lymphoblastoid cell lines with ALS-associated heterozygous hnRNPUL1 mutations A50T or S249N **(Figure S5F)**; in both cases, patients were confirmed negative for ALS-associated mutations within the major ALS risk genes SOD1, C9ORF72, TARDBP and FUS (data not shown). Levels of hnRNPUL1 were significantly reduced with the A50T mutation **(Figure S5G)**, similar to the levels within the hnRNPUL1-AID cells in the absence of Dox/Aux. This prompted us to test whether the defect in snRNA 3′ processing was already present in the hnRNPUL1-AID cells in the absence of Dox/Aux. Measurement of the ratio of the downstream region, past the cleavage site, to the snRNA gene body via qRT-PCR revealed the processing defect was readily detected in the hnRNPUL1-AID cells in the absence of Dox/Aux **(Figure 5G)**. Since the hnRNPUL1-AID cells have normal CB levels in the absence of Dox/Aux, this indicates that the processing defect is direct, rather than an indirect effect via perturbation of CBs. Analysis of the SPRY/dPNK AlphaFold prediction revealed that the S249N patient mutation could cause protein misfolding around the SPRY region through loss of stabilising hydrogen bonds between neighbouring flexible strands, potentially disrupting the overall structure **(Figure S5H)**. The reduced levels of hnRNPUL1 in the A50T cells and potential protein folding disruption caused by the S249N mutation both suggested potential loss of hnRNPUL1 function and consistent with this, we observed a processing defect on U4 snRNA in these cells **(Figure 5H)**. We focused on U4 snRNA as it was the most sensitive to hnRNPUL1 loss out of all tested snRNAs **(Figure S5A)**. Since ALS leads to motor neuron death, we generated motor neurons differentiated from human pluripotent stem cells containing an AID-tagged hnRNPUL1 gene **(Figures S5I and S5J)** and again observed the U4 snRNA processing defect, indicating this phenotype is present in disease-relevant cells **(Figure 5H)**. We were able to complement the snRNA processing defect in the HCT116-derived hnRNPUL1-AID cell line by stable expression of WT, but not ΔWA or ΔRGG mutants **(Figures 5I and S5K)**, indicating that nucleotide triphosphate binding, together with RNA binding, are essential for its function in snRNA 3′ processing. Despite our best efforts, a ΔWB-expressing cell line could not be generated for complementation analysis, despite facile transient expression **(Figure 2C)**, suggesting stable expression of the construct was toxic.

### hnRNPUL1 is required for histone transcription and pre-mRNA cleavage

snRNAs play diverse roles in the cell beyond splicing and U7 is required for cleavage and maturation of nascent histone pre-mRNAs. This, together with the significant enrichment of hnRNPUL1 on U7 snRNA **(Figure 3A)**, prompted us to analyse the effects of hnRNPUL1 loss on histone mRNA processing. Northern blotting of U7 snRNA showed decreased levels relative to the PolIII-transcribed U6 snRNA following hnRNPUL1 depletion **(Figure 5J)**. Mammalian native elongating transcript sequencing (mNET-seq) revealed decreased replication-dependent histone gene transcription upon loss of hnRNPUL1, which may result from a direct role in transcription or feedback from poor termination **(Figure 5K)**. Although the ChIP-seq binding profile over histone genes showed strong hnRNPUL1 enrichment over promoters **(Figure S5L)**, its highest enrichment was observed downstream of the annotated transcript end sites **(Figure 5L)**, suggestive of involvement in regulating the downstream cleavage of nascent histone pre-mRNAs. To monitor the extent of histone misprocessing within our HCT116-derived hnRNPUL1-AID, we performed poly(A)-selected nuclear RNA-seq, which captures polyadenylated misprocessed histone pre-mRNAs. Depletion of hnRNPUL1 led to an increase in polyadenylated histone mRNAs, whilst mature histone mRNAs, observed in our sRNA-seq, were depleted, confirming cells required hnRNPUL1 for their proper processing **(Figure 5M)**. In support of these data, differential expression analysis of public K562 hnRNPUL1 siRNA RNA-seq datasets showed an increase in poly(A)-selected histone levels, revealing these termination defects are observable within other systems **(Figure S5M)**.

### hnRNPUL1 recognizes the terminal stem loop of snRNAs and phenocopies INTS11 depletion

Our data indicate hnRNPUL1 is required for PolII-mediated snRNA 3′ end cleavage and transcription termination, which is governed by the Integrator complex **(Figures 5D, 5F and S5C)**. To rule out the possibility that hnRNPUL1 loss simply prevents Integrator:PolII interactions, we analysed their association in the presence and absence of hnRNPUL1 **(Figure 6A)**. The levels of subunits from every Integrator module^58^ associated with PolI remained unchanged following hnRNPUL1 depletion, demonstrating that hnRNPUL1 is unnecessary for Integrator binding to PolII. Next, we investigated whether there was a direct interaction between hnRNPUL1 and Integrator in cells transfected with WT and ΔWA variants **(Figure 6B)**. Co-IP revealed an interaction between Integrator (represented by the endonuclease INTS11 and PolII phosphatase PPP2AC) and both WT and ΔWA, consistent with an earlier report of hnRNPUL1 binding INTS6^59^. The observation that hnRNPUL1:Integrator:PolII complex association was resistant to ribonuclease suggests that direct protein interactions mediate their association.

**Figure 6.**
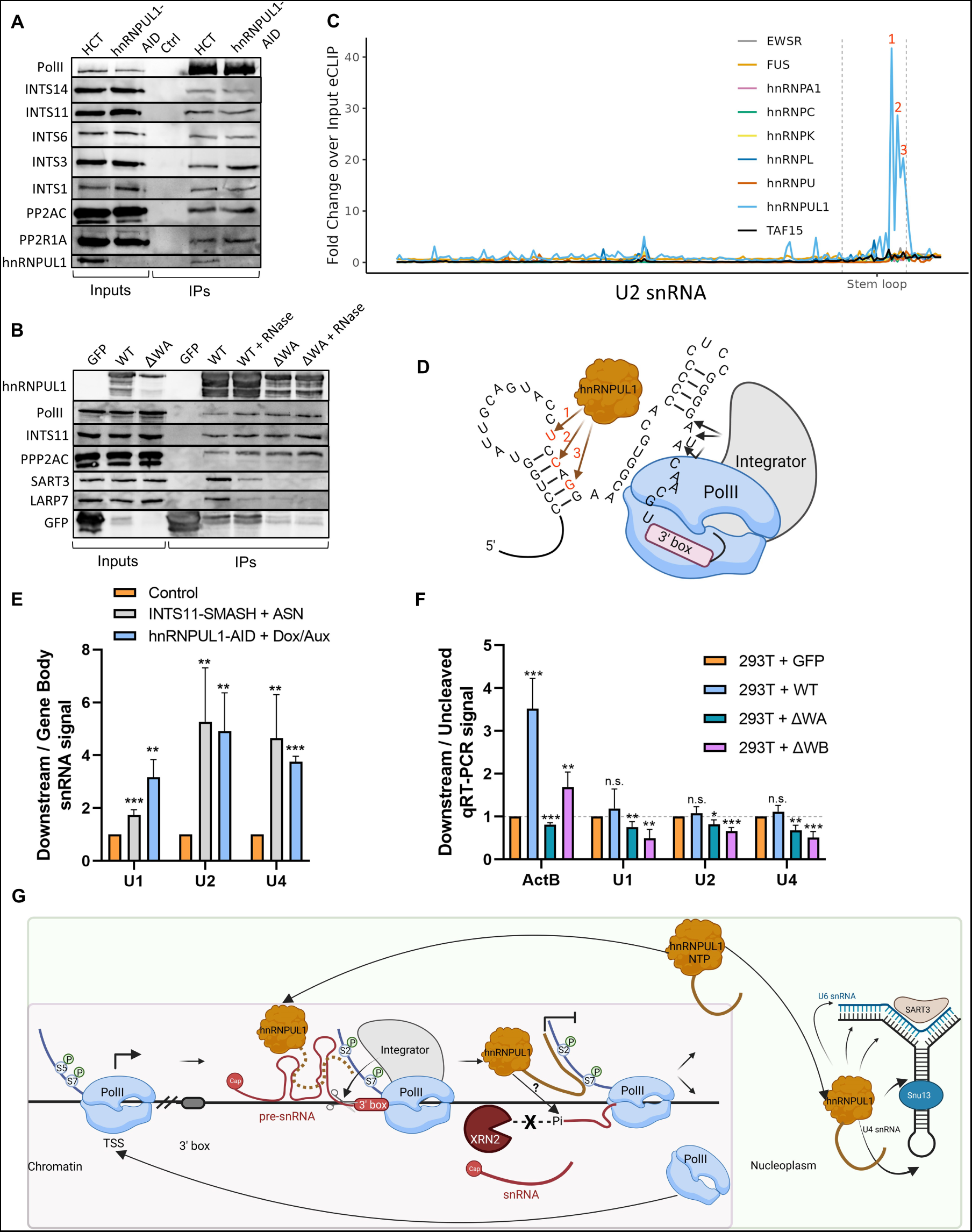
hnRNPUL1 binds the Integrator complex and facilitates its action. **(A)** PolII co-IP with components of the Integrator complex in hnRNPUL1-AID cells. **(B)** hnRNPUL1 WT/ΔWA co-IP with PolII and components of the Integrator complex before and after RNase A digestion. FLAG-tagged hnRNPUL1 variants were transfected in HEK293T cells. SART3 and LARP7 are shown as RNase-sensitive controls. **(C)** eCLIP signal over the U2 snRNA from selected hnRNPs. **(D)** Diagram of U2 snRNA 3′ region highlighting the position of the hnRNPUL1 peaks from (A) (numbered and coloured red) and the cleavage site for Integrator. **(E)** Levels of 3′-extended snRNAs measured using the ratio between downstream and gene body amplified snRNA levels in ASN treated INTS11-SMASH cells, relative to untreated cells (n=4); and hnRNPUL1-AID cells relative to Dox/Aux-treated HCT116 cells (n=3). **(F)** qRT-PCR ratio of downstream/uncleaved signal at specified loci after overexpression of hnRNPUL1 WT/ΔWA/ΔWB by transient transfection in HEK293T cells (n=4). The bar charts show relative mean values from at least three independent experiments, with the error bars showing s.d. All statistical analyses were performed using two-tailed t-tests. n.s. = not significant, *p<0.05, **p<0.01, ***p<0.001. **(G)** Model for hnRNPUL1 activity at snRNA loci and in di-snRNP recycling. The brown dotted/solid line indicates the flexible hnRNPUL1 CTD which includes the RGG domain potentially disrupting secondary structure in RNA substrates.

To understand how hnRNPUL1 might assist Integrator in the cleavage process, we revisited hnRNPUL1 eCLIP and focused on the localised enrichment over snRNAs, with other hnRNPs included as controls. hnRNPUL1 alone exhibited a ∼40-fold enrichment within the 3′ end of the U2 snRNA **(Figures 6C and 6D)**, in a stem loop previously shown to be required for the accuracy of Integrator cleavage^60^. We also observed enrichment within the terminal stem loops of the U7 and U11 snRNAs, suggesting hnRNPUL1 binds upstream of the 3′ processing site on snRNAs to stimulate cleavage **(Figures S6A and S6B)**. To put the 3′ processing defect elicited by hnRNPUL1 loss in context, we compared the effects of loss of the integrator subunit INTS11 or hnRNPUL1 on cleavage of nascent PolII-transcribed snRNAs. INTS11 was depleted from modified HCT116 INTS11-SMASH cells by treatment with asunaprevir (ASN), as reported previously^51^. Strikingly, a comparable 2-5-fold increase in unprocessed U1, U2 and U4 snRNA transcripts was observed following both INTS11 and hnRNPUL1 depletion **(Figure 6E)**, revealing the vital role that hnRNPUL1 plays in Integrator-mediated cleavage of snRNAs.

### hnRNPUL1 overexpression can stabilise an XRN2 substrate

Whilst the 5′-mono-Pi RNA fragment generated by cleavage of protein coding mRNAs is degraded by XRN2, ultimately leading to transcription termination, this is not the case for the equivalent RNA fragment generated following snRNA and histone 3′ processing. How these fragments evade XRN2 remains unresolved, given it ChIPs downstream of both snRNAs and histones^61^. Instead, these downstream cleavage products appear to be degraded from the 3′ end by the exosome via the NEXT complex^62^, which is detected in our MS analysis of hnRNPUL1 interactors. Co-IP analysis confirmed these interactions, including a particularly strong RBM7 interaction **(Figure S6C)**. We considered that hnRNPUL1 might utilise its dPNK domain to bind such 5′-mono-Pi RNA species generated by INTS11 and protect them from XRN2, thus dictating an alternative transcription termination pathway. Unfortunately, these RNA species are highly transient, leaving us unable to detect binding to them either in existing eCLIP data or via RIP. However, we noticed that hnRNPUL1 had a robust ChIP signal downstream of protein coding genes, most notably *ActB* **(Figure S6D)**. To test the capacity of hnRNPUL1 to protect 5′-mono-Pi RNA from XRN2, we examined the impact of hnRNPUL1 WT and mutant overexpression on the RNA level downstream of the *ActB* cleavage/polyadenylation site, which is an XRN2 substrate **(Figures 6F, S6E and S6F)**. We used the ratio of downstream/uncleaved RNA to normalise for effects on transcription. The overexpression of WT led to increased levels of the downstream RNA, consistent with inhibition of XRN2 activity. In contrast, ΔWA, which shows reduced RNA binding to a mono-Pi RNA via the dPNK domain, and is trapped in the chromatin fraction, failed to increase levels of the XRN2 substrate. ΔWB, which shows enhanced RNA binding within the dPNK domain **(Figure 1E)**, but is also trapped on chromatin, did increase the downstream RNA levels, albeit not as efficiently as wild-type. The response of U1, U2 and U4 snRNAs to hnRNPUL1 overexpression contrasted with that on *ActB*. The wild-type protein had no significant effect on the ratio of downstream to uncleaved RNA, indicating it is not limiting at these loci, whereas ΔWB had a dominant negative phenotype, significantly increasing the uncleaved RNA levels, but not the downstream levels **(Figures 6F, S6E and S6F)**. Together, these data indicate that increased levels of WT and to a lesser extent ΔWB hnRNPUL1 can disrupt XRN2-mediated degradation of the ActB RNA downstream of the 3′ cleavage site, whereas snRNA genes are insensitive to increased WT levels, and ΔWB overexpression inhibits their 3′ cleavage.

## Discussion

We have identified hnRNPUL1 as a component of the snRNA 3′ cleavage machinery, which associates with the PolII-Integrator complex. It binds a terminal hairpin in the U2 snRNA, which was previously implicated in 3′ processing fidelity of this snRNA^60^ and in a *Xenopus* extract this sequence was necessary for 3′ cleavage^63^. In a purified system, Integrator cleaves substrates with hairpin secondary structures poorly and also without obvious specificity^20^, suggesting that *in vivo* other factors may licence this cleavage reaction, including hnRNPUL1. The hnRNPUL1 binding site in U2 snRNA is centred around 25 bases upstream from the Integrator cleavage site, which intriguingly is similar to the typical distance between a polyadenylation site and the cleavage site in a pre-mRNA. hnRNPUL1 has an RGG RNA binding domain, which in the case of FUS, destabilises base stacking, unfolding an RNA loop^64^. Thus, the hnRNPUL1 RGG domain, which is essential for activity **(Figure 5I)**, may serve a similar function to prepare the substrate for INTS11 cleavage, or alternatively use this region to bind the terminal stem loop in snRNAs, positioning hnRNPUL1 close to the Integrator cleavage site **(Figure 6F)**. Depletion of hnRNPUL1 also reduced PolII levels over the snRNA gene body, which may be through a direct role in snRNA transcription initiation and elongation or negative feedback mechanisms due to an accumulation of PolII in the termination region **(Figure 5F)**. We also observed a defect in histone mRNA 3′ processing, which in part can be attributed to reduction in the levels of U7 snRNA. However, hnRNPUL1 is a prominent U7 snRNA interactor **(Figure 3A)**, which may indicate a role beyond that in U7 biogenesis, with the mature U7 snRNP as suggested previously^18^.

The RNA downstream of the 3′ processing site on mRNAs is degraded by XRN2, which recognises the 5′-monoPi RNA, ultimately catching up with PolII and causing transcription termination. How XRN2 cleavage of RNAs might be regulated is poorly understood. For example, the downstream cleavage products of INTS11, CPSF73 and DROSHA at snRNA, histone mRNA and pri-miRNA genes, respectively, are not degraded by XRN2 and do not rely on XRN2 for transcription termination^51, 61, 65–67^. In hnRNPUL1, we have identified a novel abundant nuclear 5′-mono-Pi RNA binder, utilising a dead polynucleotide kinase domain which has the potential to compete with XRN2 for substrates, thus dictating both degradation and transcription termination pathways. Whether the related proteins hnRNPU and hnRNPUL2 play a similar role potentially on different RNA substrates remains to be determined. hnRNPUL1 has also been implicated in repair of DNA DSBs by homologous recombination, though the mechanism was unclear^15^. DSBs lead to bidirectional transcription of dilncRNAs, which are subsequently processed by DROSHA, and again, the termination of these transcription events does not involve XRN2^67^. Intriguingly, hnRNPUL1 was identified as a prominent Microprocessor complex interactor^68^, suggesting hnRNPUL1 might have a direct role in transcription termination and degradation pathways for dilncRNAs, coupled with DROSHA binding.

The evolutionary preservation of the potential to form an active polynucleotide kinase by mutation of a single amino acid in hnRNPUL1 is remarkable **(Figure 1K)**, indicating that many of the original functions of the ancestral fold have been preserved, including nucleotide and nucleic acid binding and the conformational changes associated with binding and release of substrates. These properties now appear to be harnessed for reversible binding to RNA and allowing hnRNPUL1 to operate in both the chromatin and nucleoplasmic environments, where hnRNPUL1 takes part in distinct processes in snRNP biogenesis and recycling. Since snRNA processing and recycling take part in CBs, it is striking that they are strictly dependent on hnRNPUL1 for formation. However, this phenotype is common to BRAT1, another Integrator co-factor^69^. BRAT1 mutations lead to neurodevelopmental and neurodegenerative disorders, whereas we identified hnRNPUL1 mutations in ALS patients. However, ALS leads to death of motor neurons, in common with SMA, which involves mutations in SMN, required for snRNP maturation^70^. Notably, hnRNPUL1 loss also disrupts the formation of Gems marked by SMN. This indicates substantial overlap in their functions, leading to a common disease basis for ALS and SMA as suggested previously for FUS, a prominent hnRNPUL1 and SMN interactor whose mutation in ALS also prevents the formation of Gems^54^. A recent report identified mutations in hnRNPUL1 associated with patients with congenital limb malformations, a phenotype quite distinct from ALS, yet these patients also had PODXL, CFTR and PRKD2 gene mutations, which could be acting as modifiers altering disease outcome^16^.

In conclusion, we propose that hnRNPUL1 is an essential cofactor for Integrator-mediated cleavage of snRNAs **(Figure 6G)**. It utilises an unstructured CTD to interact with PolII CTD, ensuring it is available to interact with nascent snRNA as it leaves the PolII active site and stimulate its timely cleavage by Integrator. Subsequently, the cleaved 3′ RNA fragment may be bound by the dPNK domain, forcing an exosome-mediated degradation pathway and XRN2-independent termination mechanism. Nucleotide triphosphate binding induces a conformational change and allows reversible binding of RNA, allowing hnRNPUL1’s reuse in the two compartments where it operates. In the nucleoplasm, it drives recycling of the U4:U6 di-snRNP together with SART3. The strong preference for U4 snRNA is most likely driven by the interaction with Snu13, which nucleates other components of the snRNP^41^, while the RNA contacts between hnRNPUL1 and U4/U6 snRNA are probably governed by RNA binding regions other than the dPNK domain, which is unable to accommodate a capped RNA. The Snu13 interaction might also explain why hnRNPUL1 is enriched on box CD snoRNAs **(Figure 3B)**, which also contain Snu13, and H/ACA snoRNAs via interaction with the Snu13-related protein NHP2. The robust RNA-mediated interactions between hnRNPUL1 bound primarily to U4, and SART3 bound to U6 snRNA no doubt help drive the efficient reassembly of the U4/U6 di-snRNP for further rounds of splicing.

## METHODS

### Lead contact

Further communication and requests for resource sharing should be addressed to the lead contact Stuart A. Wilson.

### Materials availability

New reagents generated in this study are available via the lead contact.

### Data and code availability

All raw sequencing datasets from this study have been submitted to the NCBI Gene Expression Omnibus (GEO) under accession number GSE228810. Custom code used in this work is available at: https://github.com/sudlab/yonchev_et_al.

### Experimental model and study participant details

HEK293T, HCT116 cell lines were maintained in Dulbecco’s modified Eagle medium (DMEM) with 10% fetal bovine serum (FBS). FLAG-hnRNPUL1 cell lines were generated using the Flp-IN T-Rex system by expression of pcDNA5-FRT/FLAG-hnRNPUL1 and pOG44 Flp recombinase expression vectors in Flp-In T-Rex 293 cells, followed by 100 μg/ml Zeocin and 15 μg/ml Blasticidin antibiotic selection. Cells were maintained in 100 μg/ml Hygromycin following selection and FLAG-tagged protein expression was induced using tetracycline addition. Lymphoblastoid cell lines were maintained in RPMI 1640 medium supplemented with 20% FCS and 200 μl L-glutamine.

#### Bacterial culture

*E. coli* strains were grown at 37°C in LB medium for plasmid propagation and in TB medium (12 g/l tryptone, 24 g/l yeast extract, 12.54 g/l K_2_HPO_4_, 2.31 g/l KH_2_PO_4_, 4 ml/l glycerol) for protein overexpression.

#### Generation of auxin-inducible hnRNPUL1 degron

HCT116 cells were subjected to CRISPR/Cas9 genome editing to tag hnRNPUL1 with an AID. Cells were seeded in a 6-cm dish, and 24 hrs later transfected with 1 µg of Cas9-expressing guide RNA plasmid targeting the *hnRNPUL1* locus, along with 1 µg of two repair plasmids containing either hygromycin or neomycin selection markers. A P2A sequence that is cleaved upon translation separated selection markers from AID. After 48 hrs of transfection, cells were split in a 10-cm dish with 150 µg/ml hygromycin and 800 µg/ml neomycin. Selection was maintained until single colonies grew, which were transferred to individual wells in 24-well plate and expanded for western blot screening. Expression of osTIR1 in homozygous hnRNPUL1-AID cells was achieved via CRISPR/Cas9-mediated insertion of TIR1 gene in safe-harbour locus AAVS1^71^. The OsTIR1 expression vector comprised a TIR1 gene controlled by a conditional Tet promoter and a puromycin resistance gene. The repair template consisted of AAVS1 locus homology arms flanking this region. hnRNPUL1-AID cells were seeded in a 6-cm dish and after 24 hrs transfected with 1.5 µg of osTIR1 vector and 1.5 µg of Cas9-expressing plasmid with AAVS1 locus-targeted guide RNA. Cells were split in a 10-cm dish 48 hrs after transfection and selected by addition of 1 µg/ml puromycin. Cells were cultured until individual colonies formed, which were transferred to individual wells in a 24-well plate and expanded. Integration of TIR and activation of the AID system were screened by treatment with 1 μg/ml doxycycline for 72 hrs and 500 μM auxin for 24 hrs. Cells were harvested and hnRNPUL1 depletion was assessed via western blot. Successful colonies were used in time courses to identify minimum doxycycline and auxin incubation periods required for full hnRNPUL1 depletion. As a result, 48 hr doxycycline and 2 hr auxin treatments were performed in subsequent experiments unless otherwise stated.

#### Generation of SB hnRNPUL1-AID

hnRNPUL1-AID lines constitutively expressing WT or mutant hnRNPUL1 were generated using the optimised sleeping beauty (SB) transposon system^72^. hnRNPUL1-AID cells were seeded in a 6-cm dish and transfected with 1.5 µg SB plasmid containing *hnRNPUL1* and 500 ng pCMV(CAT)T7-SB100. SB plasmid (derived from pSBbi-blast) contained a blasticidin resistance gene for selection of hnRNPUL1 integration. Cells were split in a 10-cm dish 48 hrs after transfection and treated with 20 µg/mL blasticidin until individual colonies formed, which were screened for hnRNPUL1 expression by Western blot.

#### Generation of hnRNPUL1-AID hPSC

Stable lines with AID-tagged hnRNPUL1 were generated by growing human embryonic stem cell line H7 (WA07)^73^ in E8 on vitronectin and at passage number p2+4+8+7 (each ’+’ denotes defrost) dissociating and transfecting 1.5 x 10^6^ cells with 1.5 µg of each px330 hnRNPUL1 gRNA, hnRNPUL1 AID-Hyg and hnRNPUL1 AID-neo with Neon microporator kit (1600 V, 20ms, 1pulse). Cells were replated onto vitronectin in E8 with 10 µM Y-27632. G418 (50 µg/ml) and hygromycin (25 µg/ml) were added to the medium 24 hr post-transfection. G418 and hygromycin concentrations were gradually increased over several days to final concentrations of 125 µg/ml G418 and 50µg/ml hygromycin in E8. Colonies were picked and expanded in mTESR1 on geltrex. For constitutive TIR1 expression, one clone was transfected with SB-TIR1 and transposase using a Neon microporator kit as described above, before selection in 10 µg/ml blasticidin.

#### hPSC differentiation to motor neurons

hPSC (3,000 cells) were plated on day 0 in 96 well U-bottomed low attachment plates in N2B27 media (50:50 neurobasal and DMEM F-12, N2 1:100, B27 1:50, Glutamax 1:100, non-essential amino acids 1:100, Beta-mercaptoethanol 1:1000) containing 5 μM Y-27632 (only for plating), 0.2 μM LDN193189, 3 μM Chir99021, 40 μM SB431542 and 0.05% PVA. Plates were spun at 400x *g* for 4 min. Media was changed every 2 days and only half media was replaced each time. On day 2, 0.1 μM retinoic acid and 0.5 μM SAG were added to the media. On day 7, LDN, Chir and SB were removed from the medium and 10 ng/ml BDNF and 10 ng/ml GDNF were added. On day 9, EBs were pooled and dissociated for 30 min using accutase in a shaker set to 37°C and 800-1400 rpm. Speeds increased by 200 rpm every 5 min after EBs were manually pipetted up and down. Dispersed cells were replated at 52,000 cells/cm^2^ in media with 10 μM DAPT, 10 ng/ml BDNF, 10 ng/ml GDNF, 0.1 μM retinoic acid and 0.5 μM SAG. Plates were prepared with Poly-L-Ornithine for 30 min at 37°C, washed 3x with PBS and coated with Geltrex (1:100). Half the media was changed every three days. On day 14, retinoic acid and SAG were removed and DAPT increased to 20 μM. On day 16, 10 ng/ml CNTF was added. On day 17, DAPT was removed. Subsequently, media contained only BDNF, GDNF, CNTF at 10 ng/ml. Cells were used for experiments on day 33.

#### Generation of FlpIN FLAG-hnRNPUL1 Stable Cell Lines

FlpIn-293 (10,000 cells) were seeded in a 6-cm dish and cultured in Tet-free FCS DMEM for 24 hrs at 37°C before transfection with 2.4 μg FRT vector and 3.6 μg FlpIn recombinase (pPGKFLPobpA) using Turbofect transfection reagent. Cells were split into two 10-cm dishes 48 hrs later. Selection medium (DMEM containing 15 μg/ml Blasticidin and 0.1 mg/ml Hygromycin) was supplied until individual colonies grew, which were transferred to wells in 24-well plates and expanded to allow screening via western blotting.

## METHOD DETAILS

### Plasmids

Cloning was performed using Gibson assembly with a kit from NEB. Point mutations were introduced with an NEB site-directed mutagenesis kit. Plasmids were checked by Sanger sequencing.

### Transfection

Cells were seeded 24 hrs before transfection to achieve ∼70% confluency at the time of transfection. Lipofectamine 2000 was used to transfect cells at 5 μl / 2.5 μg of DNA in a mixture of serum-free DMEM for a final volume of 1:10 to the dish volume. Cells were harvested 48 hr later. For protein overexpression in mammalian cells, HEK293T were seeded in 10-cm dishes, similarly. Cells were transfected with a mixture of PEI:DNA of 3.5:1 (w/w) in serum-free DMEM for a final volume of 1:6 to the dish volume and harvested 48 hr later.

### Cell fixation and Immunostaining

Cells grown on coverslips were washed in 1x PBS and incubated in fixing solution (4% formaldehyde, 1x PBS) for 20 min. Cells were washed and incubated in permeabilisation solution (0.5% Triton-X 100, 1x PBS) for 10 min, then incubated in 1% BSA solution with primary antibody for 1 hr, washed 3x in 1x PBS, and incubated in 1% BSA with secondary antibody for 30 min. Following three 1x PBS washes, coverslips were mounted onto slides using VECTASHIELD DAPI-containing medium.

### Immunoblotting

Western blotting was performed on Amersham Protran nitrocellulose membrane within a Trans-Blot Turbo Transfer System. Proteins were transferred at 25 V, 1.3 mA for 22 min. Blots were incubated in TBST + 5% milk for 1 hr, then with primary antibody for 1 hr, washed in 1x TBST 3x 5 s and 3x 5 min. Incubation (1 hr) with secondary antibody (at 1:10,000) coupled to horseradish peroxidase (HRP) followed. Membranes were washed as before, placed in ECL detection reagent for 1 min and exposed in a Bio-Rad Chemidoc System.

### Recombinant protein purification

#### FLAG-tagged proteins

HEK293T cells were lysed in 5 pellet volumes IP Lysis buffer (IPLB) (50 mM HEPES-NaOH pH 7.5, 100 mM NaCl, 0.5% Triton-X 100, 1 mM EDTA pH 8.0, 10% Glycerol, 1 mM DTT, EDTA-free Protease Inhibitors) and nuclei were sheared by aspiration with a 26-gauge needle. hnRNPUL1 ΔWA lysates were digested with 500 U benzonase at 37°C for 30 min. Lysates were centrifuged at 16,200x *g* for 5 min and supernatants transferred onto 100 µl FLAG agarose beads pre-washed with IPLB. Beads were incubated at 4°C for 2 hr, washed with 2x1 ml IPLB and treated with 4 µg RNase A at 37°C for 30 min. Beads were washed with 3x1 ml IP wash buffer (IPLB + 1 M NaCl) and once with 1 ml IPLB. Proteins were eluted in 300 µl IPLB + 0.25 µg/µl 3xFLAG peptide at 4°C.

#### Hexahistidine-tagged proteins

*E. coli* BL21 (DE3) cells were resuspended in 10 pellet volumes Co^2+^ lysis buffer (50 mM Na_3_PO_4_ pH 8.0, 300 mM NaCl, 10% Glycerol, 0.5% Triton-X 100) and sonicated (10x [30s-ON/30s-OFF]) at 80% amplitude with a Fisherbrand Model 120 Sonic Dismembrator. Lysate was centrifuged at 20,000x *g* and 4°C for 30 min. Supernatants were treated with 5 μg RNase A for 20 min at 37°C, then loaded onto a 5 ml self-packed Talon Superflow Cobalt column. Beads were washed with 1 column volume (CV) Co^2+^ lysis buffer, then 1 CV lysis buffer + 5 μg RNase A and incubated for 30 min. The column was washed with 6 CV Co^2+^ wash buffer (Lysis buffer + 10 mM Imidazole) and the protein eluted with 2 CV Co^2+^ elution buffer (50 mM Na_3_PO_4_ pH 7.0, 300 mM NaCl, 150 mM Imidazole). Proteins were buffer-exchanged into Co^2+^ elution buffer without Imidazole by repeated centrifugation in Vivaspin 10K MWCO protein concentrators.

### RNA oligonucleotide labelling

An RNA oligonucleotide with 5′ hydroxyl was end-labelled with T4 PNK. The reaction contained 0.5 μM RNA, 2 μl T4 PNK 10X buffer (NEB), 16.7 μM ^32^P γ-ATP, 10 U T4 PNK (NEB) and was incubated at 37°C for 1 hr, then RNA was separated on a 10% denaturing polyacrylamide gel in 0.5x TBE buffer (0.44 M Tris, 0.44 M Boric Acid, 1 mM EDTA, pH 8.0). The RNA was cut from the gel and crushed, then extracted overnight with 400 μl RNA gel extraction buffer (1 M NaCH_3_COO^-^, 1 mM EDTA). The supernatant was filtered through a Spin-X centrifuge tube and precipitated with 1 ml 100% ethanol and 5 μg glycogen. After 1 hr at -20°C, the RNA was centrifuged at 16,200x *g* for 30 min. The pellet was washed with 1 ml 75% ethanol, centrifuged at 16,200x *g* for 7 min, air dried and resuspended in 100 μl nuclease-free water.

### NTP crosslinking and competition assays

#### ATP UV-crosslink

Proteins (0.3 μM) were mixed in NTP binding buffer (NTP RB: 50 mM Tris-HCl pH 8.0, 100 mM NaCl, 10 mM MgCl_2_, 1 mM DTT) and 55 nM ^32^P γ-ATP. Reactions were incubated on ice for 15 min and UV-crosslinked on ice for 30 min at 254 nm. Crosslinked proteins were separated on an 8% gel by SDS-PAGE, stained and dried at 80°C for 30 min. Dried gels were exposed on a phosphorimager screen, which was developed on a TyphoonFLA 7000 laser scanner.

#### NTP competition

Full-length hnRNPUL1 (1.6 μM) was mixed in NTP RB with 55 nM ^32^P γ-ATP and AMP-PNP or GMP-PNP non-hydrolysable competitors (final concentrations: 0.55, 5.5 and 27.7 μM). Reactions were mixed for 15 min on ice and UV-crosslinked for 30 min. Complexes were separated on a 10% gel by SDS-PAGE and processed as before.

#### GTP/m^7^G cap analog competition

hnRNPUL1 ΔCTD (1 μM) was mixed in NTP RB with 10 nM α-^32^P GTP and GTP or m^7^G cap analog competitors at 0.1, 1 and 10 μM. Samples were incubated at 37°C for 15 min and UV-crosslinked on ice for 30 min. Complexes were separated on a 10 % gel by SDS-PAGE and processed as before.

### RNA UV-crosslinking and competition

#### RNA UV-crosslink

SPRY/dPNK proteins (0.5 μM) were mixed in RNA binding buffer (15 mM Hepes pH 7.9, 100 mM NaCl, 5 mM MgCl_2_, 0.2 mM EDTA, 0.5 % Tween-20, 10 % glycerol) with ∼25 nM 5′-labelled RNA oligonucleotide and 8 U RNAse inhibitor and incubated on ice for 10 min, then UV-crosslinked on ice for 15 min. Full-length proteins were additionally incubated with 10 mM ATP in the reaction mixture. Subsequently, proteins were separated on a 10% gel by SDS-PAGE and processed as before.

#### ATP/ADP competition

hnRNPUL1 ΔCTD (0.5 μM) was incubated in NTP RB with ∼25 nM 5′-labelled RNA oligonucleotide, 8 U RNase inhibitor and ATP or ADP competitors at 10, 100 and 1,000 μM. Reactions were incubated at 37°C for 10 min and UV-crosslinked on ice for 15 min. Complexes were separated on a 10% gel by SDS-PAGE and processed as before.

### Kinase assays and <u>T</u>hin <u>L</u>ayer <u>C</u>hromatography (TLC)

#### Kinase assays

Proteins (0.5 μM) were mixed in NTP RB with 0.5 μM dsDNA/RNA oligonucleotides with 5′ hydroxyls and 15 nM ^32^P γ-ATP. The T4 PNK control contained 10 U enzyme and 2 μl 10X T4 PNK buffer instead. Reactions were incubated at 37°C for 30 min and separated on a 12% 6 M urea polyacrylamide gel in 0.5X TBE. Gel was exposed on a phosphorimager screen for 1 hr and developed as before.

#### TLC

TP hydrolysis reactions were performed in NTP RB with 0.5 μM protein, 0.8 nM ^32^P γ-ATP and 0.5 μM RNA oligonucleotide (as specified). Reactions were incubated at 37°C for 30 min and quenched with 0.25 M EDTA and 10 μl xylene cyanol. Proteinase K (20 μg) and 100 mM Tris pH 7.5 were added and proteins were digested for 30 min at 37°C. T4 PNK and apyrase positive controls were performed differently. T4 PNK reactions contained 5 μl 10X T4 reaction buffer, +/- 0.5 μM RNA oligonucleotide, 0.8 nM ^32^P γ-ATP and 10 U T4 PNK. The apyrase control contained 5 μg apyrase, 5 μl 10X NTP binding buffer and 0.8 nM ^32^P γ-ATP. Reactions were incubated at 37°C for 30 min, then quenched with 0.25 M EDTA and 20 μl xylene cyanol. PEI-cellulose plates were pre-run in 0.4 M phosphate buffer pH 5.5 and dried. Equal amounts of radioactivity were spotted on PEI plates and run in the same buffer, dried and exposed on a phosphorimager screen for 1 hr and developed as before.

### Fluorescence spectroscopy studies

SPRY/dPNK proteins were buffer-exchanged into 50 mM Tris pH 8.0, 100 mM NaCl by serial concentration/dilution with Pierce protein concentrators to dilute the 3XFLAG peptide used for elution to 0.025 ng/μl. Proteins (0.5 μM) were mixed with 50 μl 10x Trp Fluo Buffer (0.5 M Tris pH 8.0, 1 M NaCl, 0.1 M MgCl_2_, 50 mM DTT) in 0.5 ml total volume. Tryptophan fluorescence was measured with a Cary Eclipse fluorometer (Agilent), excitation wavelength= 280 nm and emission spectra= 300-400 nm with spectral resolution of 5 nm and photomultiplier set to high. Reaction buffer + 0.025 ng/μl 3XFLAG peptide was used to determine minimal background fluorescence and subtracted from subsequent measurements. Each protein was measured in apo form, then ATP was titrated into the solution. Final ATP concentrations tested were: 10, 25, 50, 74, 99, 246, 491, 735, 978, 2430 and 4840 nM. After each titration, proteins were incubated for 1 min at 37°C and emission spectra recorded in triplicate. Five values around the emission peak (347-352 nm) were averaged for each spectrum and fluorescence quenching (ΔFluorescence) was calculated for each set of triplicate measurements. Curves were fitted in Graphpad Prism 9 corresponding to a One site– Specific binding equation.

### RNA extraction and qRT-PCR

Cells grown in 6-cm dishes were washed with 1x PBS and lysed in 250 μl of IPLB containing protease inhibitors and RNase inhibitors. Trizol LS (750 μl) was added for 10 min at RT. Chloroform (200 μl/750 μl TRIzol) was added and samples were shaken, then centrifuged at 12,000x *g* for 15 min at 4°C. The top layer was transferred to a new tube and RNA precipitated with 1:1 isopropanol, centrifugation at 12,000x *g* and washing with 75% EtOH. Resulting pellets were air dried and resuspended in 43 μl dH_2_O. RNA was incubated with Turbo DNase for 1 hr at 37°C and precipitated using RNA acidic phenol pH 5.5, 85 mM NaAc pH 5.8 and 70% EtOH. Pellets were washed twice in 75% ethanol and resuspended in RNase-free water. Total RNA (1 μg) was reverse-transcribed into cDNA with High Capacity cDNA Reverse Transcription Kit. cDNA was diluted 10x and 4 μl were mixed with 5 μl 2x Sensimix and 1 μl 5 μM primer mix. qRT-PCR was performed on a Corbett Rotor-Gene 2000 instrument for 45 cycles.

### Cellular Fractionation

Cells were trypsinized and transferred to 15 ml Falcons in pre-warmed DMEM-FCS, then spun down at 400x *g* for 3 min and washed twice in 1x PBS. Pellets were transferred to 2 ml round-bottom Eppendorfs and resuspended in 7x volumes of ice-cold sucrose lysis buffer (10 mM Tris-HCl, pH 8.0, 0.5 M Sucrose, 10% glycerol, 3 mM CaCl_2_, 2 mM MgCl_2_) with 0.5% Triton-X 100, 1 mM DTT, EDTA-free Protease and RNAse Inhibitors for 5 min, before spinning at 500x *g* for 5 min at 4°C. The supernatant was centrifuged at 16,000x *g* and kept as cytoplasmic fraction, with 250 μl added to 750 μl Trizol-LS for RNA extraction, and 50 μl kept for western blotting. Nuclear pellets were washed twice in sucrose lysis buffer, once in 1x PBS, and once in sucrose lysis buffer. Pellets were gently resuspended in 300 μl NRB (20 mM HEPES, pH 7.5, 75 mM NaCl, 1 mM DTT, 50% Glycerol, EDTA-free Protease Inhibitors) and 300 μl NUN buffer (20 mM HEPES, pH 7.5, 300 mM NaCl, 1 mM DTT, 10 mM MgCl_2_, 1 M Urea, 1% NP-40) were added for 5 min invering every minute. Samples were spun at 1,200x *g* for 5 min at 4°C. The supernatant was centrifuged at 16,000x *g* and kept as nuclear fraction, with 250 μl added to 750 μl Trizol-LS for RNA extraction, and 50 μl kept for western blotting. Pellets were resuspended in 1 ml buffer A (10 mM HEPES, pH 7.5, 10 mM KCl, 10% glycerol, 4 mM MgCl_2_, 1 mM DTT, EDTA-free Protease Inhibitors), transferred to 1.5 ml Eppendorf and spun for 5 min at 1,200x *g* and 4°C. The supernatant was removed and the pellet resuspended in 250 μl buffer A and added to 750 μl Trizol-LS for RNA extraction. Pellets were resuspended in 100 μl RIPA buffer and incubated with 1 μl benzonase (125 U/μl) for 45 min at RT for protein extraction. The reaction was centrifuged at 16,000x *g* for 10 min at 4°C and 50 μl supernatant was kept for western blotting.

### RNase/DNase release assay

Cellular fractionation was performed on two 6-cm dishes as described above until collection of a chromatin pellet in Buffer A. The pellet was washed twice and centrifuged at 1,200x *g* for 5 min at 4°C, then resuspended in 100 μl Buffer A with 0.25 mg/ml RNase A, and incubated at 37°C for 15 min. After centrifugation at 1,200x *g* for 5 min, the supernatant was collected as the RNase-soluble fraction. The cell pellet was washed twice with 1 ml Buffer A, resuspended in 100 μl Buffer A with 1X Turbo DNase buffer and 10 U Turbo DNase, and incubated at 37°C for 30 min. After centrifugation at 16,000x *g* for 5 min, the supernatant was collected as the DNase-soluble fraction.

### Immunoprecipitation

#### Formaldehyde RNA immunoprecipitation (RIP)

For one 6-cm dish/RIP, cells were at 80% confluency for experiment. Protein-G Dynabeads (100 μl/condition) were washed twice in RIP lysis buffer (RIPLB: 50 mM HEPES-HCl pH 7.5, 150 mM NaCl, 0.1% SDS, 1% NP-40, 0.5% sodium deoxycholate, 10% glycerol, EDTA-free Protease Inhibitors) before resuspension in 400 μl RIPLB + 1% BSA and 4-10 μg of relevant antibody with rotation for 1 hr. Beads were then washed 3x in RIPLB before addition of sample lysate. Protein-RNA complexes were crosslinked through incubation with PBS-formaldehyde (0.1%) for 10 min, before quenching with addition of glycine (0.125 M final concentration). Cells were washed 3x with PBS and lysed in RIPLB + Ribosafe RNase Inhibitors and Turbo DNase. Sonication using a Bioruptor (5 x [30s-ON/30s-OFF]) generated RNA fragments of 300-400 nucleotides. Samples were cleared by centrifugation (16100x *g*, 15 min, 4 °C), and 10% total lysate was isolated for the input fraction. Remainders were loaded on beads and IPs performed whilst rotating for 2 hrs at 4 °C. This was followed by washing the beads twice with RIPLB, then twice with RIP high salt wash buffer (50 mM HEPES-HCl, pH 7.5, 1 M NaCl, 0.1% SDS, 1% NP-40, 0.5% sodium deoxycholate, 10% glycerol), and finally two more washes with RIPLB. Input and IP samples were then made up to 56 μl with ultrapure H_2_O, followed by the addition of 33 μl 3X reverse crosslinking buffer (3X PBS, 6% N-lauroyl sarcosine, 30 mM EDTA, pH 8, 15 mM DTT), 190 μl proteinase K (19 mg/ml) and 1 μl Ribosafe RNase Inhibitors. Samples were incubated for 1 hour at 42 °C and then 1 hour at 55°C, both whilst shaking at 1100 rpm, to reverse the crosslinks and digest proteins in the samples. RNA was then isolated from input and IP samples via TRIzol extraction (detailed above) and converted into cDNA prior to qPCR analysis.

#### Chromatin immunoprecipitation (ChIP)

For each ChIP condition, 1-4 15-cm dishes were seeded with 5x10^6^ cells/dish. Protein-DNA complexes were crosslinked through incubation with 20 ml PBS-formaldehyde (1%). Cell pellets were lysed in ChIP Lysis Buffer 1 (50 mM HEPES-NaOH pH7.5, 140 mM NaCl, 1 mM EDTA, 0.5% NP-40, 0.25% Triton-X 100, 10% glycerol, Protease Inhibitors) and rotated for 5 min at 4°C. Nuclei were pelleted by centrifugation (3000x *g*, 5 min at 4°C), resuspended in ChIP Buffer 2 (10 mM Tris-HCl, pH 7.3, 200 mM NaCl, 1 mM EDTA, 0.5 mM EGTA, Protease Inhibitors) and rotated for 10 min. Nuclei were pelleted by centrifugation (1500x *g*, 5 min at 4°C) and resuspended in ChIP Lysis Buffer 3 (10 mM Tris-HCl pH7.3, 200 mM NaCl, 1 mM EDTA, 0.5 mM EGTA, 0.5% N-lauroylsarcosine, 0.1% Na-deoxycholate, Protease Inhibitors). Chromatin fragments of 250-300 nucleotides were generated by sonication in a Diagenode Bioruptor pico (20 x [30s-ON/30s-OFF]). Samples were cleared by centrifugation (16100x *g*, 15 min, 4°C) and equal concentrations of chromatin were incorporated into IPs, performed overnight at 4°C using 5 μg antibody. Blocked protein-G Dynabeads (100 μl) were added to samples and incubated for 2 hrs at 4°C, then beads were washed 4x with ChIP RIPA Wash Buffer (50 mM HEPES-NaOH, pH 7.5, 500 mM LiCl, 1% NP40, 1 mM EDTA, 0.1% N-lauroylsarcosine, 0.7% Na-deoxycholate) and once with ChIP Final Wash Buffer (10 mM Tris-HCl pH7.3, 50 mM NaCl, 1 mM EDTA). Complexes were eluted with ChIP Elution buffer (50 mM Tris-HCl pH8.0, 10 mM EDTA, 1% SDS) over 30 min at 65°C. NaCl (200 mM) was added and cross-links were reversed overnight at 65 °C. Samples were treated with RNase A (0.2 mg/ml final) for 2 hrs at 37°C, followed by proteinase K (0.2 mg/ml final) for 2 hrs at 55°C. DNA was purified via phenol-chloroform extraction and ethanol precipitation, and resuspended in H_2_O.

#### Protein co-ImmunoPrecipitation (co-IP)

Dynabeads Protein-G (40 μl) were blocked by rotating at 24 rpm for 2 hrs in 400 μl IPLB + 4 μg primary antibody and 1% BSA. Beads were washed 3x in 1 ml lysis buffer (IPLB) and placed on ice. IPLB + Protease Inhibitors and 1 mM DTT was placed onto 2-4 x 15-cm dishes and cells were scraped into a 1.5 ml Eppendorf and centrifuged for 5 min at 17,000x *g*. Supernatant (1-5 mg proteins) was mixed with beads and rotated for 2 hrs at 24 rpm and 4°C. Beads were washed 3x and eluted using 50 μl 1 M ArgᐧHCl pH 3.5. Eluates were neutralised with 1.5 μl Tris.HCl pH 8.8 and analysed by western blotting, with input samples representing 0.1-0.5% of protein concentration loaded onto the beads.

#### FLAG-tagged Protein co-IP

FLAG-agarose (50 μl) was blocked by rotating overnight at 24 rpm and 4°C in IPLB + 1% BSA. Cells were lysed and supernatant loaded onto beads and incubated as described above. For RNase A-treated samples, 0.25 mg/ml RNase A was supplemented during the washes before elution. Bound complexes were eluted in 60 μl IPLB + 100 μg/ml 3xFLAG peptide by rotation at 4°C for 1 hr. Eluates were analysed by SDS-PAGE or western blotting, with input samples representing 0.1-0.5% of protein concentration loaded onto the beads.

### Pulldowns

Bacterial pellets (0.5 g) were used to extract bait proteins, while 0.1-0.2 g cells were used for the control baits (GST and 6xHis-GFP). Cells were resuspended in 1 ml GST lysis buffer (1X PBS, 0.1% Tween-20) or Co^2+^ lysis buffer and sonicated (5 x [3s-ON/3s-OFF]) at 25% amplitude with a Fisherbrand Model 120 Sonic Dismembrator. Lysates were centrifuged at 16,200x *g* for 5 min and incubated with 50-150 μl GSH or Talon beads for 30 min at 4°C. Beads were washed 3x with 1 ml GST lysis buffer or Co^2+^ wash buffer, then split into several tubes (depending on number of conditions), mixed with purified prey proteins or total cell lysates (equalised by expression of transfected hnRNPUL1 constructs) in RB100 (25 mM Hepes pH 7.5, 50 mM NaCl, 10 mM MgCl_2_, 1 mM DTT, 0.05% Triton-X 100, 10% Glycerol). Reactions were incubated at 37°C for 30 min, supernatants washed with 2x 200 μl PBS and eluted with 2 bead volumes of GST-(50 mM Tris pH 8.0, 40 mM GSH, 200 mM NaCl, 10% glycerol) or Co^2+^ elution buffers. Eluates were analysed by SDS-PAGE and western blotting.

### Mass spectrometry analysis

co-IPs were conducted as above until washing and elution steps, with 10x 15-cm dishes/condition as starting material. Protein-loaded beads were washed twice with IPLB, then twice with IPLB without glycerol and Triton-X 100. Immunoprecipitated complexes were eluted using 1 M Arg-HCl pH 3.5 prior to in-solution tryptic digestion. Mass spectrometry analysis was performed using nano-flow liquid 71 chromatography on an U3000 RSLCnano coupled to a hybrid quadrupole-orbitrap mass spectrometer (Q Exactive HF). Peptides were separated on an Easy-Spray C18 column (75 μm x 50 cm) using a 2-step gradient from 97% solvent A (0.1% formic acid in water) to 10% solvent B (0.1% formic acid in 80% acetonitrile) over 5 min, then 10% to 50% B over 75 min at 300 nl/min. The mass spectrometer was programmed for data-dependent acquisition with 10 product ion scans (resolution 30,000, automatic gain control 1e5, maximum injection time 60 ms, isolation window 1.2 Th, normalised collision energy 27, intensity threshold 3.3e4) per full MS scan (resolution 120,000, automatic gain control 1e6, maximum injection time 60 ms) with a 20 sec exclusion time. Database searching MaxQuant (version 1.5.2.8) software was used for database searching with *.raw MS data file using standard settings. Data was searched against the Homo sapiens Uniprot proteome database (taxa id: 9606, downloaded 25 November 2018, 73101 entries), using the following settings: Digestion type: trypsin; Variable modifications: Acetyl (Protein N-term); Oxidation (M); MS scan type: MS2; PSM FDR 0.01; Protein FDR 0.01; Site FDR 0.01; MS tolerance 0.2 Da; MS/MS tolerance 0.2 Da; min peptide length 7; max peptide length 4600; max mis-cleavages 2; min number of peptides 2.iBAQ was selected for label free quantification.

### Colony formation assay

Doxycycline (1 μg/ml) and Auxin (500 μM) were added to cells for 48 and 24 hrs respectively, for full hnRNPUL1 depletion in the hnRNPUL1-AID degron line upon seeding. A 6-well plate/condition was seeded at 200 cells/well. Cells were left to grow at 37°C for 14 days in 2 ml DMEM, then washed with 1X PBS, followed by staining with 1ml crystal violet/well (0.5% Giemsa powder in methanol) for 5 min. Cells were washed twice with 1 ml deionized water, then left to dry for 30 min.

### Liquid-liquid phase separation assays

Assays were performed by mixing the specified protein concentration in 50 mM Tris pH 8.0, 0.1 M NaCl and 5% PEG 8,000. Small aliquots (2 μl) of each condition were spotted onto glass slides, covered with coverslips and sealed, then incubated at RT for 1 hr. Droplets settled on the slide surface were examined with a Leica fluorescent microscope. At least 5 photos were taken from representative areas. The aliphatic alcohol 1,6-hexanediol (5%) was added at the beginning of the reaction or to pre-formed droplets as indicated.

### Dual-Luciferase assay

FLAG-Gal4-fused constructs were co-transfected with a plasmid carrying Gal4-specific promoter-driven firefly luciferase (5xGAL4-TATA-luciferase) and a plasmid carrying CMV promoter-driven human codon-optimised Renilla luciferase (pGL4.75). Transfection was optimised for equal test protein expression. Dual luciferase assays were conducted using the Dual-Glo Luciferase Assay Kit as instructed. Firefly luciferase reporter activities were normalised to co-transfected Renilla luciferase activity, and the comparative transcription activation potential of each Gal4-fused construct was calculated as the relative firefly luciferase activity ratio against the Gal4-DBD negative control construct.

### Northern blot

RNA (2 μg/sample) was resolved on an 8% 8 M urea polyacrylamide gel and transferred to N+ Hybond membrane at 0.2 A for 2 hrs in 0.5x TBE. The membrane was air-dried, cross-linked under 0.2 joules UV and blocked in 20 ml Hybridization Buffer (6x SSPE [0.9 M NaCl, 54 mM NaH_2_PO_4_, 6 mM EDTA, pH 8.0], 5x Denhardt’s [0.1% (w/v) Ficoll, 0.1% (w/v) Polyvinylpyrrolidone, 0.1% (w/v) bovine serum albumin], 0.2% SDS) for 1 hr at 37°C. Oligonucleotide probe (0.25 μM) was labelled with 16.5 nM ^32^P γ-ATP and T4 PNK for 30 min at 37°C. Labelled probe was diluted with 1 ml Hybridization Buffer, denatured for 5 min at 65°C, and diluted to 5 ml with Hybridization Buffer. After filtration through a 20 μm filter, the probe was added to the blocked membrane and hybridised for 2-16 hrs at 37°C, then the membrane was rinsed 3x and washed in 6x SSPE for 1 hr at 37°C. The membrane was dried and exposed on a phosphorimager screen for 3-16 hours and visualised as before. Boiling stripping buffer (0.1x SSPE, 0.1% SDS) was used to remove the bound probe.

### Sequencing

#### Mammalian Native Elongation Transcript sequencing using mammalian cells (mNET-seq)

Mammalian NET-seq was performed as outlined by the authors^74^. Cells were seeded in 8x150-mm dishes at 40% confluency and grown overnight. Pan-specific CMA601 PolII antibody was bound to Dynabeads sheep anti-mouse IgG overnight by rotation at 12 rpm at 4°C. Upon reaching 80% confluency, cells were washed with ice-cold PBS and collected into 15-ml Falcon tubes. All centrifugation was carried out at 4°C, 420x *g* for 5 min for cells, 100x *g* for 2 s for Dynabeads IgG, and 16,000x *g* 15 min for chromatin and RNA precipitation steps, unless otherwise stated. Cells were spun down and resuspended in 4 ml HLB+N lysis buffer (10 mM Tris-HCl, pH 7.5, 10 mM NaCl, 2.5 mM MgCl_2_, 0.5% NP-40). After a 5 min incubation, cells were underlaid with HLB+NS (10 mM Tris-HCl, pH 7.5, 10 mM NaCl, 2.5 mM MgCl_2_, 0.5% NP-40, 10% Sucrose) and nuclei were harvested by centrifugation. Nuclei pellets were resuspended in 125 μl NUN1 (20 mM Tris-HCl, pH 7.9, 10 mM NaCl, 0.5 mM EDTA, 50% Glycerol), transferred to 1.5 ml Eppendorf tubes and chromatin pellets were obtained by a 15 min incubation in 1.2 ml NUN2 buffer (20 mM HEPES-KOH, pH 7.6, 75 mM NaCl, 7.5 mM MgCl_2_, 1% NP-40, 0.2 mM EDTA, 1 M Urea) and 10 min centrifugation. Chromatin pellets were washed in 100 μl MNase buffer (50 mM Tris-HCl, pH 7.9, 5 mM CaCl_2_ 100 ug/ml BSA) and incubated for 90 seconds in MNase buffer, supplemented with 40 gel units/μl MNase in an Eppendorf ThermoMixer at 37°C, 1,400 rpm. The reaction was then quenched with 10 μl EGTA and the insoluble material spun down at 16,000x *g*. Supernatants were pooled and rotated with PolII conjugated beads at 4°C for 1 hour. The beads were bound to a magnetic rack, washed 6 times with NET-2 buffer (50 mM Tris-HCl, pH 7.4, 300 mM NaCl, 0.05% NP-40), and resuspended in 300 μl PNKT buffer (70 mM Tris-HCl, pH 7.6, 10 mM MgCl2, 5 mM DTT, 0.1% Tween-20). 15 μl of the sample was processed separately for incorporation of ^32^P-ATP and monitoring the size distribution of digested RNA. The samples were resuspended in reactions containing PNKT, 1 U/μl PNK and 1.5 mM ATP (1 μCi/μl γ-^32^P ATP for size monitoring) and incubated at 37°C for 6 m at 1,400 rpm on a ThermoMixer. The beads were washed on a magnetic rack and resuspended in 1 ml TRIzol reagent and 200 μl chloroform. Following a 2 min incubation at RT, samples were spun at 16,000x *g* for 15 min and 500 μl of the supernatant was precipitated using 5 μg GlycoBlue co-precipitant and 500 μl isopropanol. After a 10 min incubation, samples were spun down and the pellets were dried, resuspended in 10 μl urea dye and run on a denaturing 8% 7 M urea polyacrylamide gel. The ^32^P-ATP incorporated sample was visualised on a Typhoon FLA 7000 laser scanner and the 35-100 nt region of the mNET-seq samples were cut out with a scalpel and placed into pierced 0.5 ml tubes. The gel slices were crushed by centrifugation into a 1.5 ml Eppendorf at 15,000x *g* for 1 min at RT. RNA elution buffer (1 M NaCH_3_COO^-^, 1 mM EDTA) was applied to the slices and they were incubated on a ThermoMixer at 1100x *g* for 2 hrs. Supernatant (400 μl) was spun through a Spin-X column and precipitated as previously described. The pellets were dissolved in RNase-free water and the sample size distributions were analysed on an Agilent 2100 Bioanalyzer. The Lexogen small RNA library prep kit was used in accordance with the manual to ligate a 5′ and 3′ sequencing adaptors to the RNA molecules, reverse transcribe them into cDNA libraries and amplify them using PCR.

#### TTchem-seq

TTchem-seq was performed as outlined by the authors^75^, with minor alterations. In brief, the following procedures were performed in the dark for each condition: two 15-cm dishes of 50% to 70% confluent cells were labelled with 500 nM 4SU for 5 minutes, rinsed with warm PBS twice, lysed and homogenized on-plate with 1.5 ml cold TRI-reagent. Each plate of cell lysate was collected into a pre-chilled 2 ml microfuge tube. The TRI-reagent-RNA mix was chloroform-extracted and ethanol precipitated. Each RNA pellet was resuspended in 200 μl ultrapure H_2_O, combined into one tube and acidic phenol-extracted using Phase Lock Gel Heavy tubes, followed by another ethanol precipitation. After resuspending the pellets in 200 μl ultrapure H_2_O, their concentration was measured using the Qubit RNA BR Assay Kit. For each sample, 100 μg of isolated 4SU-labelled total RNA was spiked in with 2 μg of 5 minute 1 mM 4TU-labelled BY4712 Saccharomyces cerevisiae total RNA and topped up to 100 μl with ultrapure H_2_O. The RNA mix was fragmented by addition of 20 μl of 1 M NaOH and was incubated on ice for 30 minutes. The fragmentation was quenched with 180 μl of 1 M Tris-HCl, pH 6.8 and immediately purified with acidic phenol-chloroform extraction and ethanol precipitation. Fragmented RNA was resuspended in 196 μl ultrapure H_2_O, and was biotinylated using 100 μg/ml Biotin-MTSEA XX in 1% Biotinylation buffer (833 mM Tris-HCl, pH 7.4, 83 mM EDTA, pH 8.0) and 50% DMF for 30 minutes at 25°C. The biotinylated RNA was sequentially acidic phenol-extracted three times, and ethanol-precipitated, followed by two 80% ethanol washes. The biotinylation was verified through a streptavidin-HRP conjugate dot-blot. The biotinylated RNA was then isolated using μMAC magnetic streptavidin beads, and washed twice with 55°C pre-heated Pull-down Wash Buffer (100 mM Tris-HCl, pH 7.4, 10 mM EDTA, pH 8.0, 1 M NaCl, 0.1% Tween-20). The captured RNA was eluted twice with freshly constituted 100mM DTT, acidic phenol-chloroform purified, ethanol precipitated, resuspended in 20 μl ultrapure H_2_O, and quantified using the Qubit RNA HS kit. The RNA fragment size was quality checked on an Agilent TapeStation system and prepared into TT-seq libraries using the NEBNext Ultra II Directional RNA Library Prep Kit for Illumina.

#### RNA-sequencing

All RNA-seq libraries were prepared by Novogene and sequenced in PE150 on an Illumina Novaseq 6000, except for mNET-seq, sequenced in SE50 on a Hiseq 2500. Chromatin RNA-seq libraries were prepared from chromatin-fractionated RNA and sequenced at 40 million read pairs/sample. Nuclear poly(A) RNA-seq libraries were prepared from nuclear RNA and sequenced at 30 million read pairs/sample. Small Nuclear RNA-seq libraries were prepared from 20-250 nt size-selected nuclear RNA and sequenced at 13 million read pairs/sample. mNET-seq and TTchem-seq libraries were sequenced at a read depth of 120 million read pairs/sample.

### Processing of sequencing data

#### RNA-seq data

RNA-sequencing datasets were adapter-trimmed using Cutadapt v1.9.1^76^ with parameters ‘-e 0.05 -- minimum-length 15’ (adapter sequences: mNET-seq: ‘-a TGGAATTCTCGGGTGCCAAGG’; TT-seq: ‘ - a AGATCGGAAGAGC -A AGATCGGAAGAGC‘; RNA-seq: ‘-a GATCGGAAGAGCACACGTCTGAACTCCAGTCAC -A AATGATACGGCGACCACCGAGATCTACAC’) and mapped to the GRCh38 genome and a junction database built from Ensembl 85, using STAR v2.4.2a^77^ with parameters ‘–outFilterType BySJout’. Bigwig tracks were generated using deeptools v3.5.1^78^ bamCoverage –binSize 1. Counts for DESeq2^79^ in sRNA and polyA RNA-seq datasets were generated using featureCounts v2.0.1^80^ with parameters ‘- t exon -g gene_id -a geneset.gtf -B -p -M -O -C’, where geneset.gtf contained exons for every annotated transcript within Ensembl 85. Genes with a baseMean count < 10 were removed from the final differential expression plots. Counts for DESeq2 in sample normalization and differential expression analysis for all other datasets were generated using deeptools ‘computeMatrix -bs 1 -m 1 -- averageTypeBins sum’ over a geneset generated from merged transcript entries for every gene, generated using cgat^81^ ‘gtf2gtf --method=merge-transcripts’. Splicing analysis of RNA-seq datasets was performed using rMATS^82^ v4.1.1 using default parameters.

#### mNET-seq data

In order to remove contaminating reads from small ncRNAs co-immunoprecipitated with RNA PolII, annotation sets of snRNA, snoRNA and scaRNA were obtained from the Hugo Gene Nomenclature Committee (HGNC) and a geneset of their chromosomal locations was generated from the hg38 RefSeq Curated table from UCSC table browser. The adapter-trimmed mNET-seq reads were mapped to a small ncRNA genome assembled from this geneset using bwa v0.7.17-r1188 mem^83^ with parameters ‘- k 20 -T 15 -a -M -t 1’. Unmapped reads were extracted and re-mapped against the GRCh38 genome using v2.4.2a^77^ with parameters ‘–outFilterType BySJout’. The resulting bam files were filtered to select primary alignments using samtools v1.9^84^ ‘view –b F 256’. Single nucleotide tracks were generated using deeptools with parameters ‘–Offset -1 –binSize 1’ and used to obtain a matrix of counts from all Ensembl gene annotations using deeptools computeMatrix. Matrix counts were used to obtain scaling factors for normalisation using DESeq2. Stranded bam files were generated using samtools view ‘-b -F 16’ and samtools view ‘-b -f 16’ for positive and negative stranded bam files, respectively. The DESeq2 scaling factors were reapplied to the stranded files using deeptools bamCoverage with parameters ‘– binSize 1 –exactScaling –scaleFactor *’ to generate stranded normalised tracks.

#### TTchem-seq data

TTchem-seq data was adapter-trimmed and mapped to GRCh38 using STAR, as above. Unspliced reads were selected using samtools ‘view %(infile)s | awk -v OFS="\t" ’$0 ∼ /^@/{print $0;next;} $6 !∼/N/’ ‘, strand-separated using samtools ‘view -b -f 128 -F 16’, ‘view -b -f 80’; and ‘view -b -f 144’, ‘view - b -f 64 -F 16’, for positive and negative strand respectively. Both bam files for each strand were merged using samtools merge. These were converted into normalised bigwigs using DESeq2 scaling factors obtained from a deeptools count matrix generated against Ensembl 85 gene annotations using deeptools bamCoverage with parameters ‘–binSize 1 –exactScaling –scaleFactor *’ to generate stranded, normalised tracks.

#### eCLIP data

eCLIP fastq files, already adapter-trimmed and UMI-extracted, were downloaded from the ENCODE project database. UMI identifiers were moved to the end of read names using a custom awk script to allow compatibility with umi-tools^85^. eCLIP reads were aligned to the human GRCh38 genome using STAR with parameter ‘--outFilterMultimapNmax 100’ to allow mapping to U2 snRNA. Aligned bam files were deduplicated using umi-tools dedup with parameters ‘--method=unique --random-seed 1 --spliced-is-unique’. Read 2 was extracted with samtools ‘view -F 64’ and counted using featurecounts with parameters ‘-M -O —read2pos 5’ for size factor generation via DESeq2. Per-base coverage of single-nucleotide eCLIP signal was extracted from the read 2 bam file into strand-specific bedgraph files using bedtools genomecov with parameter ‘-5’ for bigwig and matrix generation.

#### ChIP-seq data

Publicly available ChIP-seq datasets were adapter-trimmed and mapped using bwa v.0.7.17-r1188 mem and parameters ‘-k 20 -T 15 -a -M -t 1 -t 12’. MACS2 v.2.2.7.1^86^ was used to perform peakcalling with parameters ‘–nomodel –extsize 100 -B –SPMR –bdg -q 0.01 -g hs’. Fold change over input tracks were generated using macs2 ‘bdgcmp -m FE’.

### Metagene analysis

Metagene profiles were generated using deeptools computeMatrix. Matrices were loaded into R for processing and plotting, with bin values averaged across all replicates. The individual bin values of each gene were summed across all genes, and, excluding **Figure 5E**, the final profiles normalized by their sum. For **Figures S3C, 5C and 5D**, the bin values of each gene were first divided by the gene sum for “per-gene” normalization. snRNA genesets used in **Figures 5C, 5D and 5E** were extracted from the UCSC table browser GENCODE V43 knowngene table as a .bed file against a genelist of 16 non-variant PolII transcribed snRNAs obtained from the HGNC (group 849). These were filtered within each analysis, to only include snRNAs containing mapped reads. The histone geneset used in **Figure 5L** was obtained from the ncbiRefSeqCurated table using a genelist of all histone annotations from the HGNC (group 864). Replication dependent histone gene names from ref.^87^ were used to mark all replication dependent histones in all differential expression boxplot subfigures in **Figures 5 and S5**. snRNA and histone genesets used in bioinformatic analyses are provided within the github repository associated with this study. For eCLIP plots in **Figures 6 and S6**, a pseudocount was added to every bin prior to plotting to aid calculation of fold change over bases with no signal in the input samples.

### Publicly available deep-sequencing data

Data from public repositories was downloaded and processed as above. Datasets and corresponding accession numbers are provided within **Table S3**.

### Graphics generation

Structural models were analysed and molecular models rendered using UCSF ChimeraX v1.6^88^. Diagrams were drawn using BioRender.com.

## QUANTIFICATION AND STATISTICAL ANALYSIS

GraphPad Prism 9.1.0 software was used to generate graphs and statistical analyses of qRT-PCR data. Statistical significance was determined by a two-tailed unpaired t-test. Number of biological replicates and definition of error bars is indicated within figure legends, n.s. = not significant, *p < 0.05, **p < 0.01, ***p < 0.001. Statistical significance on differential expression data presented within box or violin plots was calculated using Welch’s t-test. All images shown are representative of experiments reproduced in at least two independent experiments, unless otherwise stated in figure legends.

## Acknowledgements

We thank Phil Mitchell for help throughout this project, Shona Murphy for providing the GST-PolII CTD construct, Vicky Porteous-Medley for technical assistance and Emma Thomson for helpful comments on the manuscript. SW acknowledges support from the Biotechnology and Biological Sciences Research Council UK (BBSRC) grants BB/W000172/1, BB/N005430/1 and BB/N014839/1. IDY, CVA and AW acknowledge support from the BBSRC White Rose Doctoral training programme grant BB/M011151/1. EAM acknowledges support from CONACyT-Mexico. MJD acknowledges support from the BBSRC (BB/M012166/1). IS acknowledges support from the BBSRC (BB/R007268/1).

## Authorship Contributions

IDY, CVA and LG performed the majority of the molecular and biochemical experiments and bioinformatic analysis, AL performed the Northern blot analysis, TT-seq and some bioinformatic analysis. HP performed the microscopy of Cajal bodies, IB and LB generated the hnRNPUL1-AID stem cells and differentiated motor neurons, respectively. EA-M performed molecular biology experiments. AW performed Gal4 dual luciferase reporter assays under supervision from AL. CE and MD performed the mass spectrometry analysis, JC-K and PS identified the hnRNPUL1 mutations in ALS patients and provided supervisory support to LG. SW provided reagents and advice on establishing the auxin degron system. IS supervised the bioinformatic analysis of all data. SAW conceived the study, designed experiments and supervised the work of IDY, CVA, LG and AL. IDY, CVA, LG and SAW wrote the manuscript with input from all co-authors.

## Competing interests

The authors declare no competing interests.

## Notes

### Competing Interest Statement

The authors have declared no competing interest.

