## Supplementary Figures for "hnRNPUL1 ensures efficient Integrator-mediated cleavage of snRNAs and is mutated in amyotrophic lateral sclerosis"

**Supporting Information**

**Supplementary Figures 1-6**

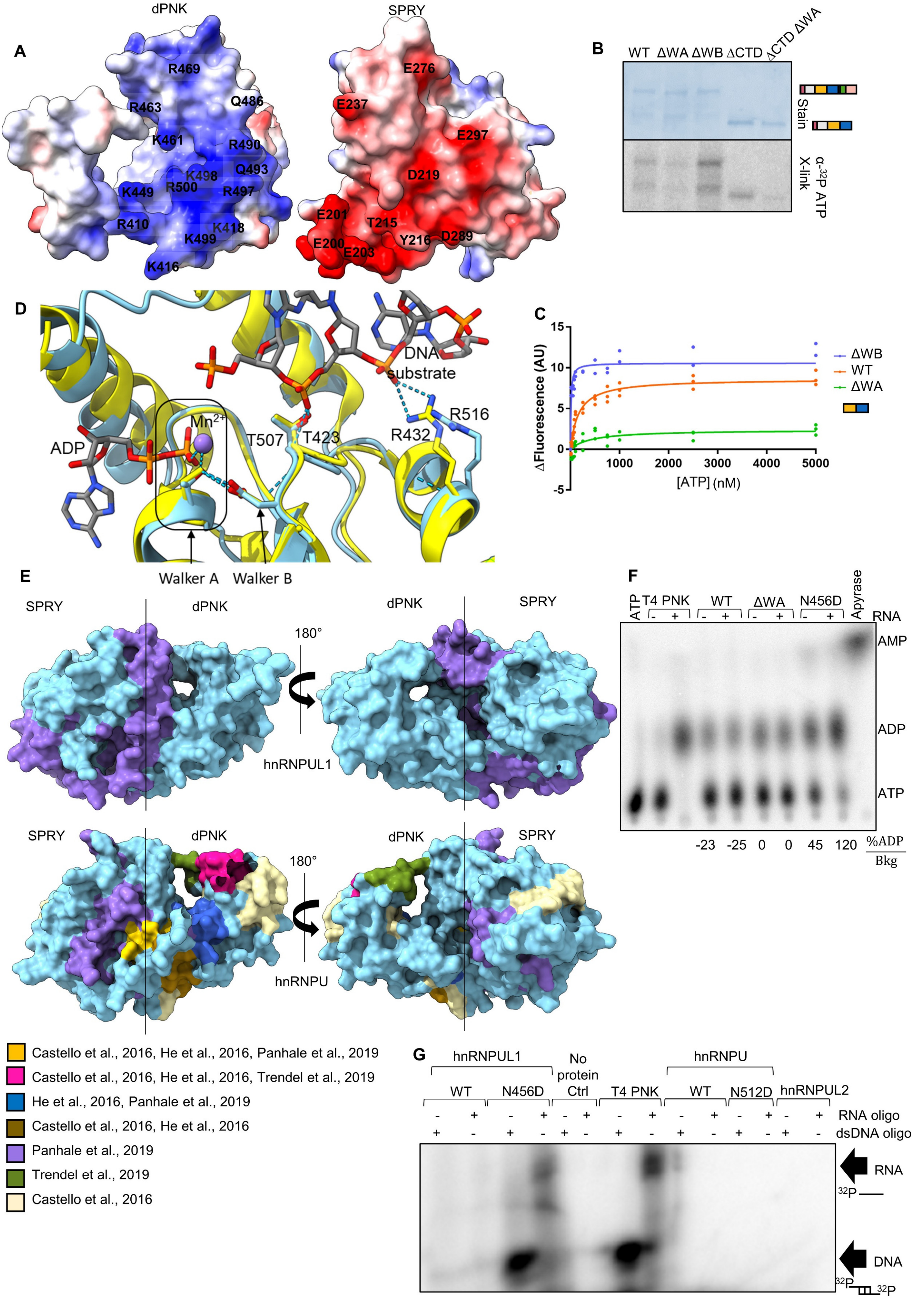


**Figure S1: hnRNPUL1 SPRY and dPNK folds form a single globular domain which binds ATP and RNA. Related to Figure 1.**

(A) Cross-section of the hnRNPUL1 core showing the interface between the SPRY and dPNK regions. Residue pairs with complementary charge are labelled.

(B) UV-crosslinking of FLAG-tagged hnRNPUL1 variants to $^{32}$P α-ATP.

(C) Tryptophan fluorescence quenching of FLAG-tagged hnRNPUL1 SPRY/dPNK core (WT= orange, ΔWA= green and ΔWB= purple) after titration of ATP.

(D) Structural comparison between hnRNPUL1 dPNK (blue) and mammalian PNKP (yellow, PDB: 3ZVN) substrate-binding pockets, highlighting ADP and DNA ligands, the WA and WB motifs and conserved residue pairs T507/T423 and R516/R432. Some regions are omitted for clarity. Dashed lines= hydrogen bonds.

(E) Surface models of hnRNPU and –UL1 cores (pale blue) with RNA-crosslinked peptides identified by mass spectrometry colour-coded based on the study that identified them.

(F) Thin-layer chromatography (TLC) tracking $^{32}$P α-ATP hydrolysis by full-length FLAG-tagged hnRNPUL1 variants. Reactions are supplemented with RNA oligonucleotide substrate from Figure 1H as indicated. T4 PNK is used as positive control for ADP generation and apyrase for AMP generation. ADP generation by ΔWA is taken as background (Bkg) for ATP hydrolysis.

(G) Kinase assay using full-length purified FLAG-tagged hnRNPU/hnRNPUL1 variants or hnRNPUL2, $^{32}$P ɣ-ATP and RNA or dsDNA oligonucleotide substrates with free 5′ ends and 5′ overhangs. T4 PNK was used as a positive control.

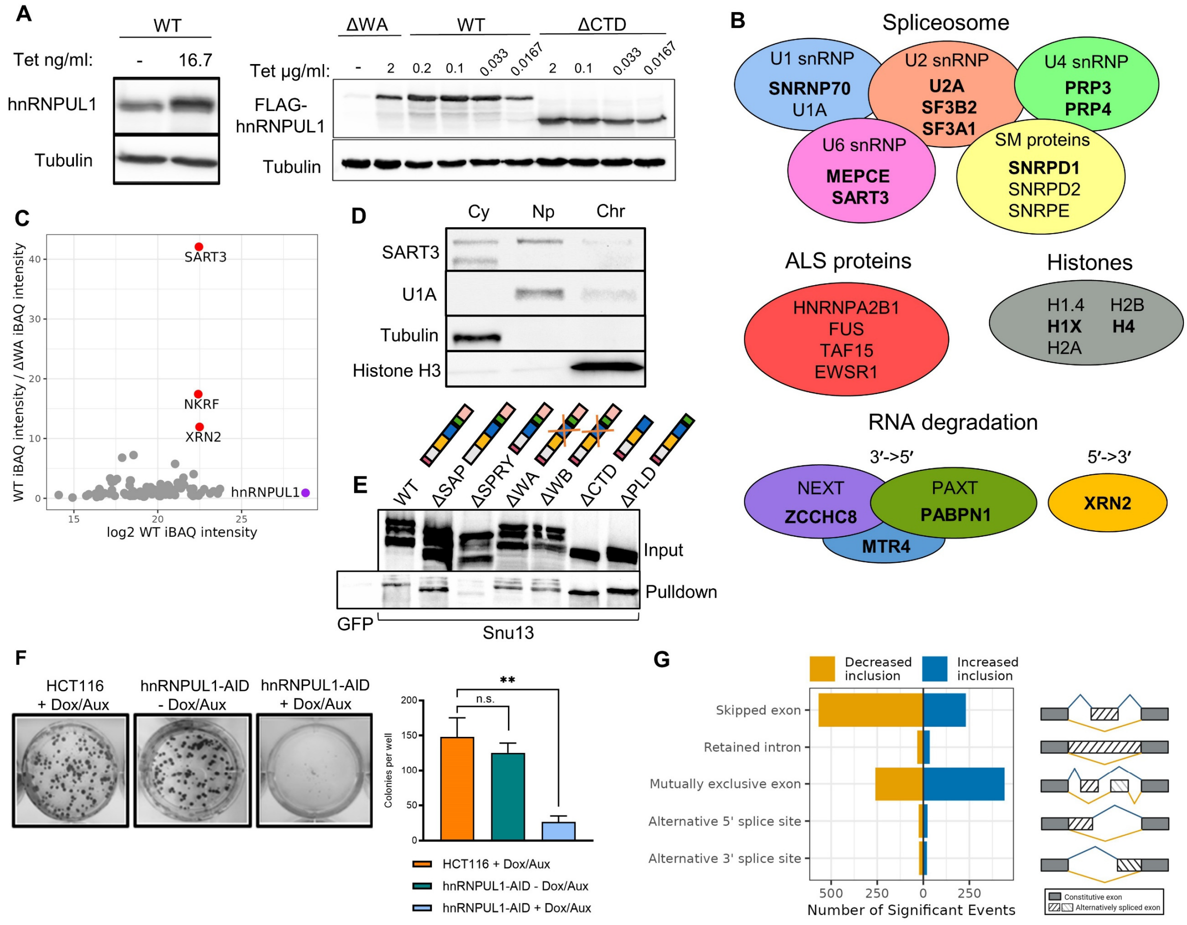


**Figure S2: hnRNPUL1 is required for U4/U6 di-snRNP formation and normal splicing. Related to Figure 2.**

(A) Expression levels of FLAG-hnRNPUL1 WT, ΔWA and ΔCTD compared to endogenous levels. Left panel = anti-endogenous hnRNPUL1 antibody. Right panel = anti-FLAG antibody.

(B) hnRNPUL1 interactors identified through Mass Spectrometry. Functionally-related proteins are indicated by coloured circles. Bold text indicates RNA-dependent interactors.

(C) Plot of Mass Spectrometry iBAQ intensity in WT condition (x axis) against the WT/ΔWA iBAQ intensity (y axis). Interactions abolished by the ΔWA mutation are highlighted in red.

(D) Fractionation of HEK293T cells. Cy= cytoplasm, Np= nucleoplasm, Chr= chromatin.

(E) Pulldowns of HEK293T lysates transfected with FLAG-tagged hnRNPUL1 variants (prey) using 6xHis-tagged Snu13 (bait) and 6xHis-tagged GFP (negative control).

(F) Colony formation assay and quantification after 14 days of treatment. The error bars show s.d. All statistical analyses performed were two-tailed t-tests. n.s. = not significant, **p<0.01.

(G) Alternative splicing events in ENCODE hnRNPUL1 siRNA dataset.

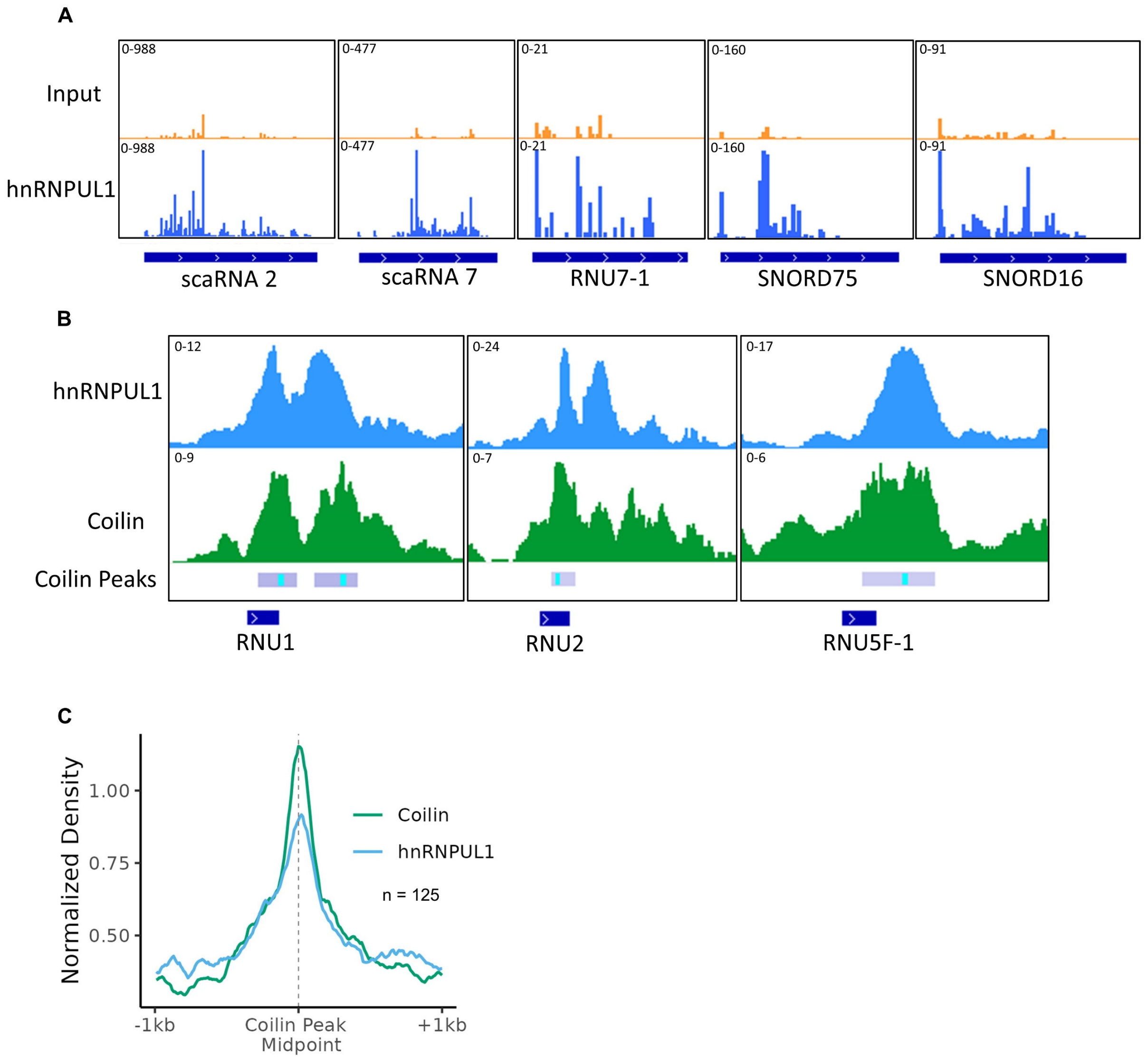


**Figure S3: hnRNPUL1 associates with noncoding RNA classes. Related to Figure 3.**

(A) IGV track screenshots of size-matched Input and hnRNPUL1 eCLIP density over two scaRNAs, two snoRNAs and U7 snRNA.

(B) IGV track screenshots of fold-change over input density of hnRNPUL1 and coilin ChIP-seq over three snRNAs. The RNU2-2P gene locus adjacent to WDR74 is shown as RNU2.

(C) Metagene of coilin and hnRNPUL1 fold-change over input density across the midpoint of 125 called coilin peaks.

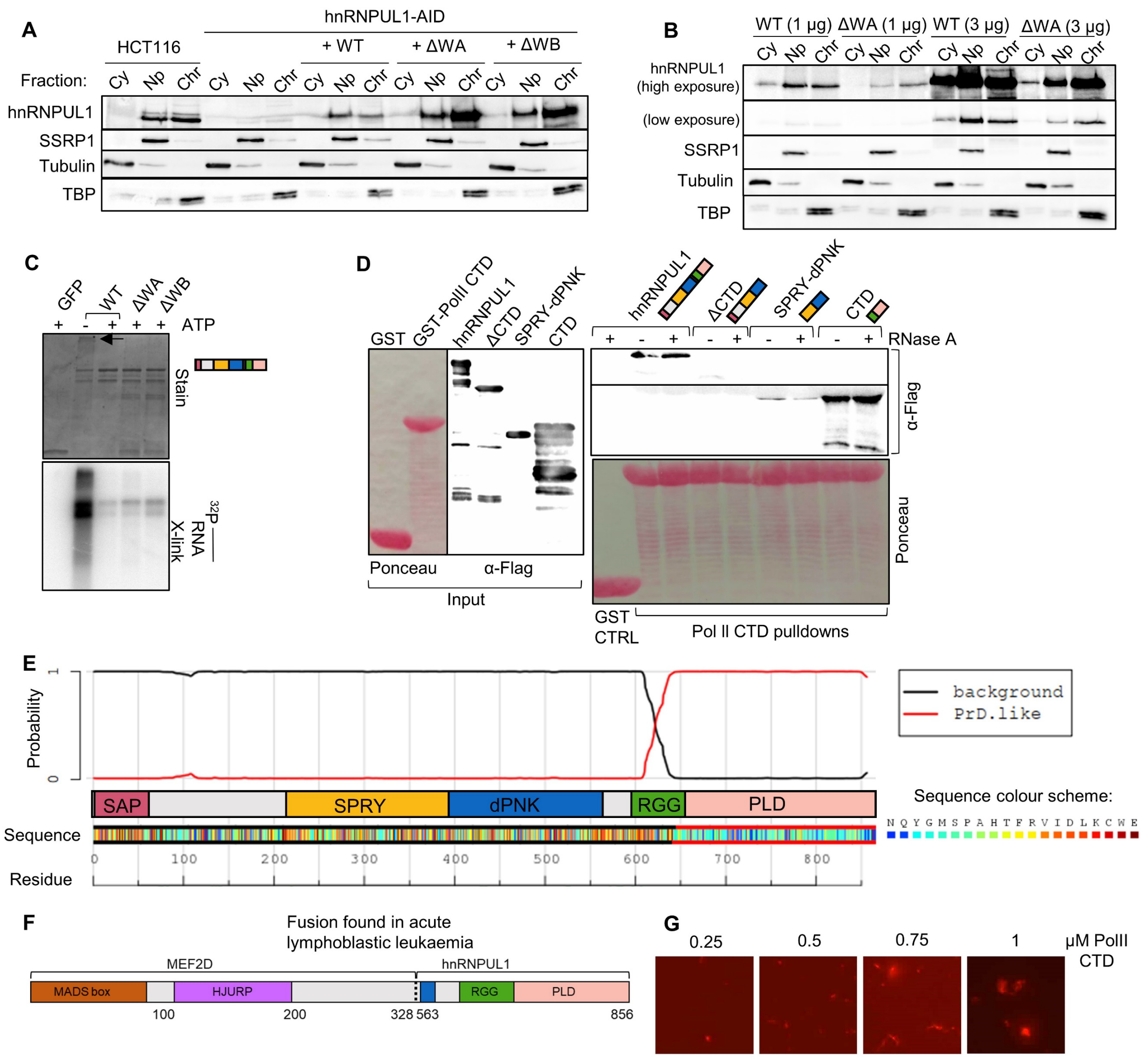


**Figure S4: hnRNPUL1 CTD mediates homo- and heterotypic protein interactions. Related to Figure 4.**

(A) Fractionation of endogenous hnRNPUL1 in HCT116 and WT/ΔWA/ΔWB hnRNPUL1 expressed in hnRNPUL1- AID cells. Cy= cytoplasm, Np= nucleoplasm, Chr= chromatin.

(B) Fractionation of FLAG-tagged hnRNPUL1 WT and ΔWA after transfection of different amounts of plasmid DNA into hnRNPUL1-AID. Cy= cytoplasm, Np= nucleoplasm, Chr= chromatin.

(C) *In vitro* UV-crosslinking of hnRNPUL1 variants to a short 5′ $^{32}$P-labelled RNA oligonucleotide in the presence of 10 mM ATP to prevent formation of protein-RNA aggregates in the wells (indicated by arrow).

(D) Pulldowns using N-terminal GST-tagged PolII CTD (bait) and N-terminal FLAG-tagged hnRNPUL1 truncations (prey). RNase A was supplemented as indicated to remove RNA contaminants.

(E) Diagram showing hnRNPUL1 amino acid composition and probability of a 60-residue region falling into the prion-like (red) or background (black) categories. Analysis was performed using PLAAC[46,47].

(F) Schematics of MEF2D-hnRNPUL1 fusion found in ALL[29].

(G) LLPS assays of PolII CTD expressed with C-terminal mCherry and N-terminal 6xHis tags at the specified concentrations.

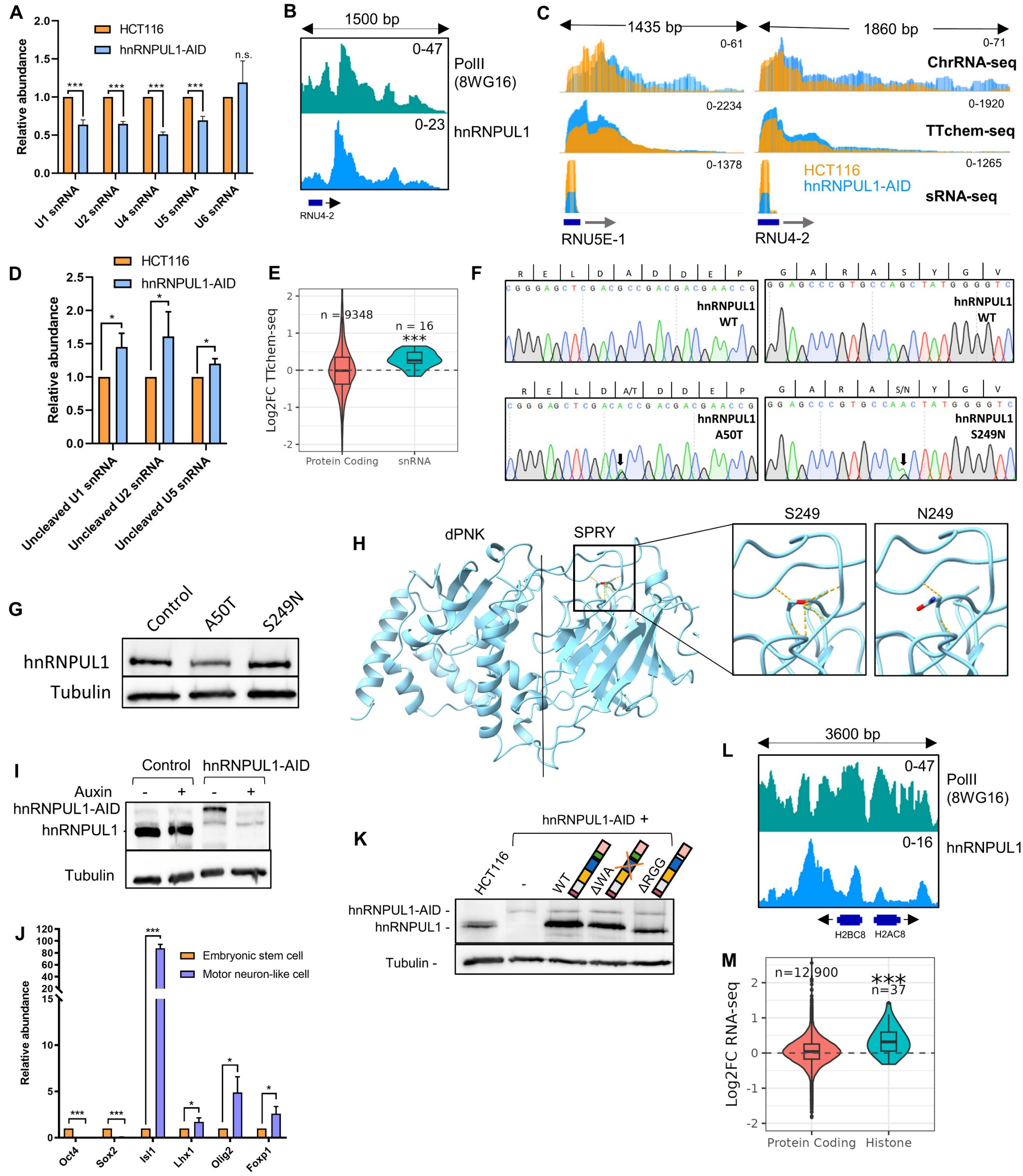


**Figure S5. hnRNPUL1 loss or ALS mutations lead to snRNA transcription termination defects. Related to Figure 5.**

(A) qRT-PCR measurement of snRNA levels, normalised to 18S rRNA, in HCT116 and hnRNPUL1 +Dox/Aux cell lines (n = 3).

(B) IGV track screenshot of PolII and hnRNPUL1 fold change over Input ChIP-seq signal over RNU4-2 and its downstream region.

(C) hnRNPUL1 chromatin (Chr), TTchem- and small RNA-seq signal over two snRNA genes and their downstream region.

(D) Uncleaved snRNA levels, measured through amplification across the 3′ box cleavage site and calculating its ratio over the total snRNA level (n = 3).

(E) Nascent protein coding and non-variant PolII-transcribed snRNA levels measured through TTchem-seq. snRNA annotations were extended 1 kilobase to accurately capture changes in uncleaved downstream fragments.

(F) Sanger sequencing trace showing heterozygosity of the A50T and S249N mutations.

(G) Protein levels of hnRNPUL1 in A50T and S249N patient lymphoblastoid cells compared to a healthy control.

(H) AlphaFold v2.0 prediction for hnRNPUL1 core highlighting changes in hydrogen bonds (orange dashes) in the S249N ALS mutation. Sidechains of S/N429 are represented in stick form and functional groups coloured by atom: red= oxygen, blue= nitrogen.

(I) hnRNPUL1 levels in motor neuron-differentiated hnRNPUL1-AID (MNhnRNPUL1-AID) human pluripotent stem cells before and after Aux treatment.

(J) qRT-PCR measurement of stem cell (Oct4, Sox2) and motor neuron (Isl1, Lhx1, Olig2, Foxp1) differentiation markers in MNhnRNPUL1-AID cells normalised to 18S rRNA (n = 3).

(K) hnRNPUL1 levels in Sleeping Beauty transposase-complemented hnRNPUL1-AID cells compared to HCT116 control.

(L) IGV track screenshot of PolII and hnRNPUL1 fold change over Input ChIP-seq signal over histone loci.

(M) Levels of replication dependent histones in ENCODE hnRNPUL1 RNAi (polyA+) RNA-seq data. Bar charts show relative mean values from at least three independent experiments with error bars showing s.d. Bar plot statistical analyses were performed using two-tailed t-tests, while box plot statistical analyses were performed using Welch’s t-test. n.s. = not significant, *$p < 0.05$, **$p<0.01$, ***$p<0.001$.

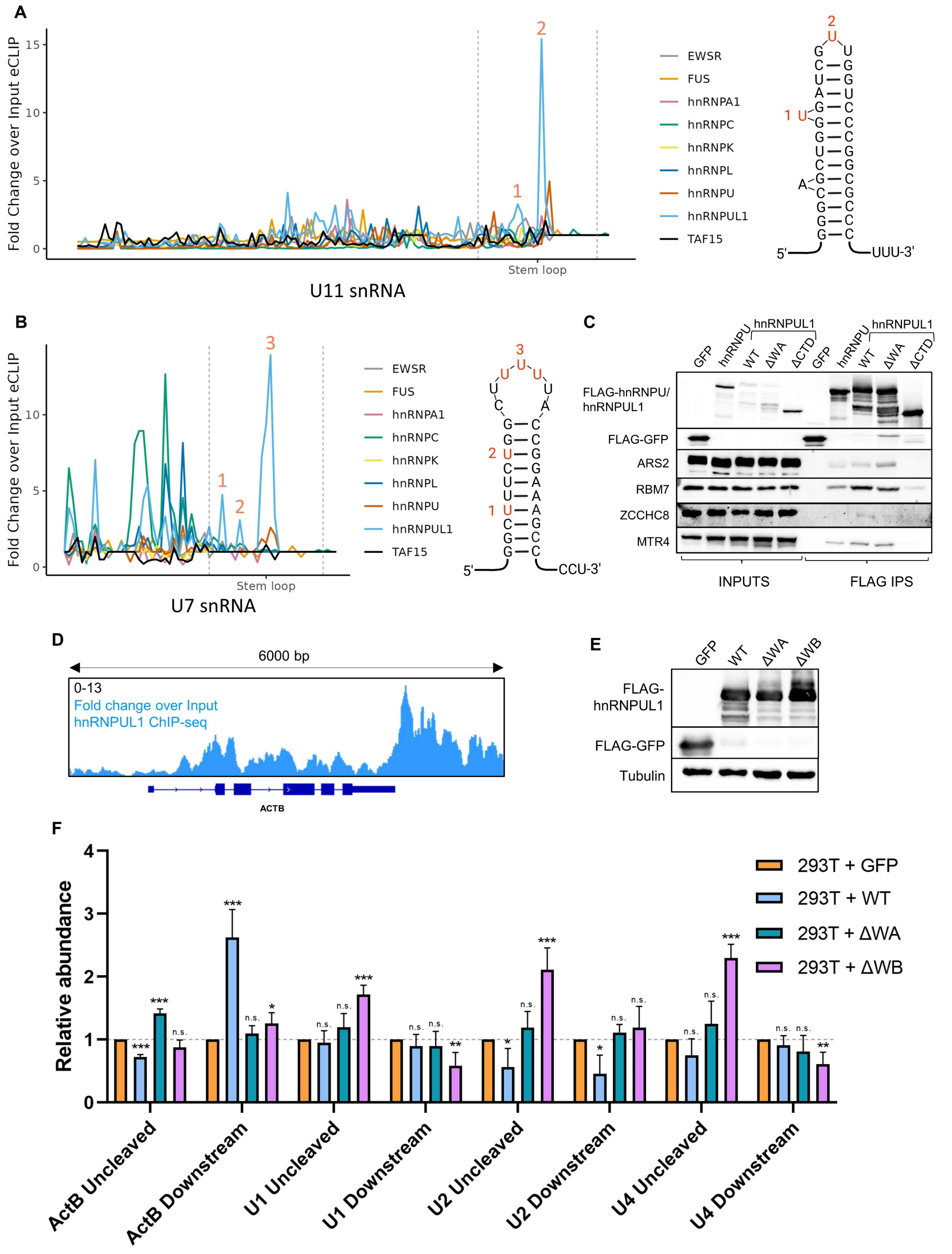


**Figure S6. hnRNPUL1 binds U snRNA terminal stem loops and can stabilise an XRN2 substrate. Related to Figure 6.**

(A) eCLIP signal over the U11 snRNA from selected hnRNPs and diagram of the terminal stem loop structure with enriched bases marked in red.

(B) eCLIP signal over the U7 snRNA from selected hnRNPs and diagram of the terminal stem loop structure with enriched bases marked in red.

(C) Co-IP of hnRNPU and hnRNPUL1 variants with members of the Nuclear Exosome Targeting Complex.

(D) IGV track screenshot of hnRNPUL1 fold change over Input signal across the ActB locus and its flanking regions.

(E) Western blot showing equal expression of FLAG-tagged proteins in HEK293T cells.

(F) qRT-PCR measurement of Uncleaved and Downstream signal at specified loci, relative to 18S rRNA, after overexpression of hnRNPUL1 WT/ΔWA/ΔWB by transient transfection in HEK293T cells (n=4). The bar charts show relative mean values from at least three independent experiments, with the error bars showing s.d. All statistical analyses were performed using two-tailed t-tests. n.s. = not significant, *p<0.05, **p<0.01, ***p<0.001.
